# Commensal skin bacteria interact with the innate immune system to promote tail regeneration in *Xenopus laevis* tadpoles

**DOI:** 10.1101/2025.05.09.653037

**Authors:** Phoebe A. Chapman, Robert C Day, Daniel T. Hudson, Joanna M. Ward, Xochitl C. Morgan, Caroline W. Beck

## Abstract

*Xenopus laevis* tadpoles regenerate their tails following partial amputation, however, for a brief developmental window, some undertake wound healing rather than regeneration. Inspired by links between microbiomes and human inflammatory disease, we asked how tadpole skin microbiomes and innate immunity influence the ability to regenerate tails. Having previously shown that lipopolysaccharide-Tlr4 signalling promotes regeneration, here, we demonstrate that the peptidoglycan-Tlr2 pathway is also involved. While tadpoles acquire their commensal skin bacteria from their mothers, analysis of mitochondrial haplotypes did not explain familial regenerative bias. Levels of endogenous lipopolysaccharides on tail tips were also not predictive of regenerative success, and shotgun sequencing indicated no difference in bacterial loads. To see if microbiome composition correlated with regenerative success, we sequenced 16S rRNA amplicons from 503 tadpole tail tips and mapped these to regenerative outcomes. While no one taxon appeared critical for regenerative success, higher proportions of Gram-positives correlated with successful regeneration. Our results suggest a previously undocumented role for Tlr2, peptidoglycan and Gram-positive commensal skin bacteria in tipping the balance from wound repair to regenerative programmes in *Xenopus laevis* refractory stage tadpoles.

## Introduction

The African clawed frog, *Xenopus laevis*, is a widely used and recognised model organism for studying development and tissue regeneration. Development from egg to metamorphosed froglet occurs outside the mother, and is well characterised into 66 stages (Nieuwkoop and Faber, 1994). Tadpoles of the species have the ability to regenerate full and functional tail tissues following partial amputation (reviewed in (Beck et al., 2009; Chen et al., 2014; Phipps et al., 2020).This is true for tadpoles at stage 40 (complete tail formation) right up to the time the tail is resorbed during metamorphosis, *except* between stages 45 and 47. In this brief “refractory” stage, tadpoles may undergo wound healing instead of regenerating lost tissues (Beck et al., 2003) providing a valuable opportunity to investigate the factors that regulate the tail amputation response. Successful regeneration depends on migration of the regeneration-organising cells (Aztekin et al., 2019) which activate regeneration-promoting secreted signalling factors such as BMP, TGFβ, FGF and Wnt (Beck et al., 2006; Beck et al., 2003; Beck et al., 2001; Chapman et al., 2022; Ho and Whitman, 2008; Lin and Slack, 2008; Sugiura et al., 2009). Tadpole tail amputation induces a rapid burst of reactive oxygen species and migration of myeloid cells towards the wound (Love et al., 2013), and these cells (likely early macrophages) are essential for the process of regeneration (Aztekin et al., 2020; Pentagna et al., 2021). Other regeneration-competent model organisms, such as zebrafish and axolotls, activate similar processes (Godwin et al., 2013; Sipka et al., 2021). Tissue-resident macrophages can be reprogrammed via their interactions with both host (damage) and microbe (pathogen) derived signals, regulating and balancing inflammation and repair processes (Chapman et al., 2022; Mosser and Edwards, 2008; Wynn et al., 2013).

The involvement of pathogen-associated molecular patterns (PAMPs) in wound healing and regeneration is becoming well documented. The microbiome has been shown to regulate responses to wounds or amputation in planaria (Arnold et al., 2016; Williams et al., 2020) and mice (Velasco et al., 2021; Wang et al., 2021). In humans, gut microbiota are linked to liver and gut regeneration (Arenas-Gómez et al., 2023; Kiseleva et al., 2024) and skin microbiota to chronic wound healing (Chapman et al., 2022; Dhankhar et al., 2024; Uberoi et al., 2024). In *X. laevis*, tail regeneration appears inherently unstable at “refractory” stages, with batch and laboratory dependent variations suggesting a stochastic process determines the dual outcomes of regeneration and wound healing. Previously, we demonstrated that tadpoles raised in antibiotics, a normal laboratory procedure for this model organism, were significantly less likely to undergo tail regeneration than their untreated siblings. We suggested that lipopolysaccharides (LPS) from Gram-negative bacteria initiate regeneration pathways by binding with Toll-like receptor 4 (Tlr4), leading to activation of the transcription factor nuclear factor κB (NF-κB) (Bishop and Beck, 2021). Indeed, regeneration could be ‘rescued’ in antibiotic-raised tadpoles by adding exogenous LPS, either commercially sourced or isolated from commensal gram negative bacteria (Chapman et al., 2022). Conversely, Tlr4 antagonistic LPS from *Rhodobacter sphaeroides* effectively suppressed tadpole tail regeneration, as did CRISPR/Cas9 knockdown of *tlr4* (Chapman et al., 2022). While the above studies have convincingly implicated LPS/Tlr4 from several microbial taxa in regeneration signalling pathways, the nature of the unstable regeneration potential in refractory tadpoles is not fully understood.

Our previous work has shown that tadpole sibships vary in both their intrinsic regenerative ability and natural skin microbiomes (Bishop and Beck, 2021; Chapman et al., 2024). Here, we ask whether the composition of the skin microbiome of tadpoles could predict their regenerative capability. In a previous study, Piccinni et al. found that adult *X. laevis* microbiomes were very similar regardless of their housing, while tadpoles were more variable and appeared more dependent on their surroundings (Piccinni et al., 2021). Compared to adult frogs, the microbiome of naturally raised refractory stage tadpoles is relatively simple, with around 65% of diversity resulting from vertical transmission from the mother frog via her egg jelly (Chapman et al., 2024). We confirm that the proportion of tadpoles undergoing regeneration varies with the mother, and that this is reproducible in subsequent spawnings in most cases. We also demonstrate a role for Tlr2 and peptidoglycan in regenerative success, but see no evidence that it is driven by either maternal relatedness or the extent of tadpole skin colonisation. 16S rRNA amplicon sequencing of 503 tail tips from tadpoles with known regeneration outcomes showed that while no individual taxon is critical for regeneration, overall higher proportions of Gram-positive bacteria on tadpole tail skin correlate with regenerative success.

## Results

### Regenerative success of a tadpole sibship varies with mother and is often reproducible between spawns

We have previously shown that tadpole sibships, when raised without antibiotics, vary widely in their predisposition to regeneration (Bishop and Beck, 2021). To see if this was inherent to the mothers, we tracked this over two separate spawns for eight females obtained from three different aquarium tanks. All eight females were induced in 2020 and then again in either 2021 or 2022. There was no difference in regenerative outcomes for 5 of the 8 groups (B4, B5, C4, D1, D3) but the remaining groups (B7, C1 and C5) had significantly different patterns of regeneration between spawns (Figure S1). Notably, these 12 sibships included very good regenerators (C4, D1, D3) where just 8 of 307 tadpoles failed to regenerate tails (2.6%) and very poor ones, like B5, where 44.8% of the 96 tadpoles failed to make any attempt to regenerate (NR category).

### Tlr2 CRISPR/Cas9 knockdown impairs regeneration while addition of peptidoglycan and/or LPS increases regenerative success

The most likely source of innate immune cells in stage 46 tadpoles are the early tissue macrophages (myeloid cells, (Aztekin et al., 2020)). We previously showed that knockdown of *tlr4* through CRISPR/Cas9 editing resulted in fewer tadpoles regenerating their tails, but that this could not account for all of the regeneration-supressing effect of raising tadpoles in antibiotics (Chapman et al., 2022). This suggests that while Tlr4 is involved in regeneration signalling pathways, additional innate immune receptors may also contribute. Another possible candidate is Toll-like receptor 2 (Tlr2), which has also demonstrated sensitivity to a range of PAMPS including peptidoglycan, an important structural component of the bacterial cell wall (de Oliviera Nascimento et al., 2012).

Interrogation of single cell sequencing data from Aztekin et al. (Aztekin et al., 2019) found that *X. laevis* Stage 46 myeloid cells express *tlr2.L* and *S* genes, along with their heterodimer partners *tlr1.L* and *tlr6.L*. We used CRISPR-Cas9 knockdown of *tlr2.L* and *S* homeologues and assayed the effect on stage 46 tail regeneration (Figure 1A, B). Partial knockdown of *tlr2* was confirmed by sequencing in three arbitrarily selected embryos per group and resulted in significant reductions in regeneration success, for two sgRNA in two different tadpole sibships (Figure 1C).

**Figure 1.**
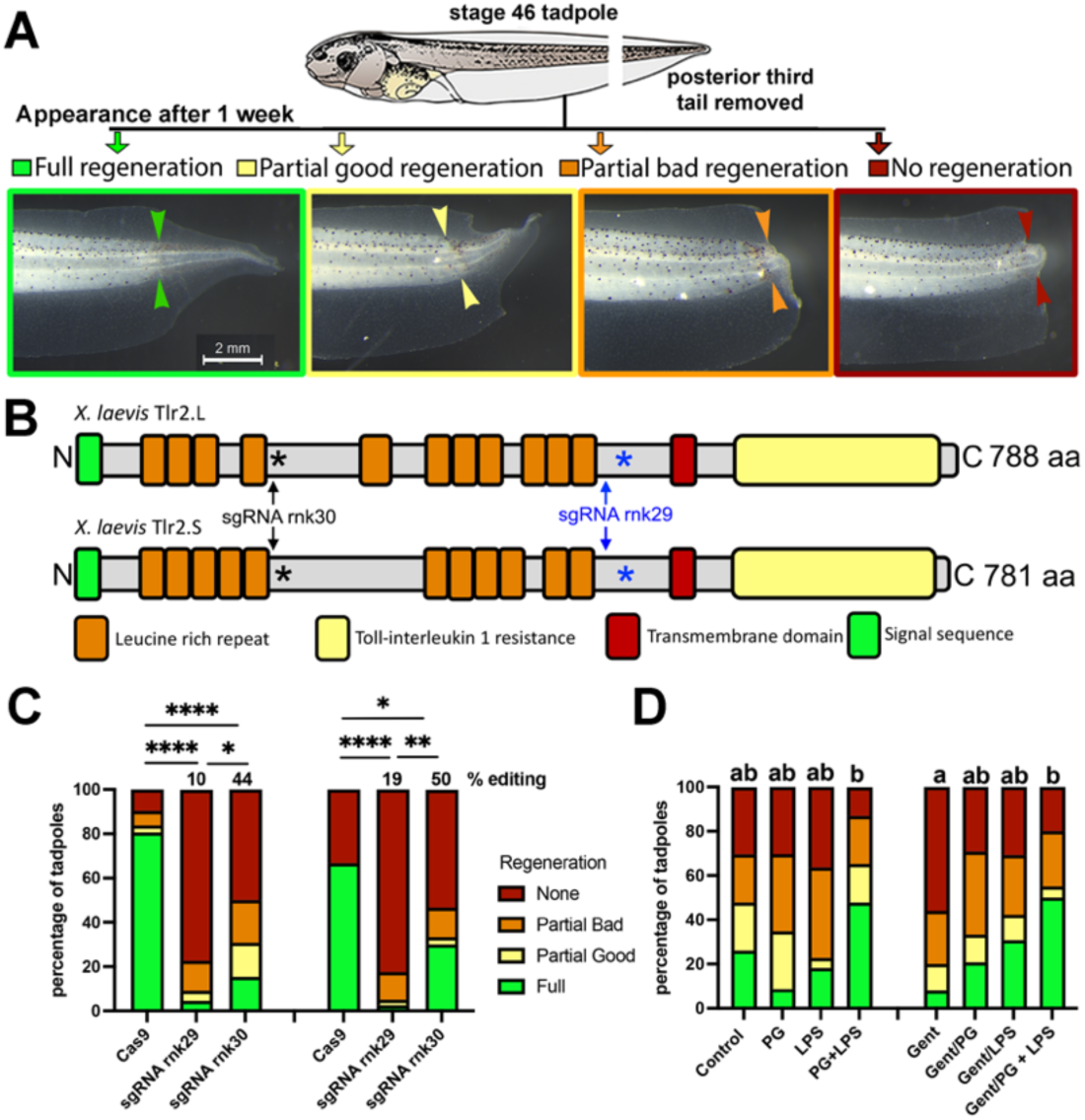
Reduced Tlr2 availability decreases the likelihood of tadpole tail regeneration. **A)** Refractory stage tail regeneration assay, with examples of the four outcomes one week after amputation. Full regeneration involves replacement of all tissues, partial good regeneration often has a minor defect such as a notch in the fin, partial bad is where one or two tissues are absent and no regeneration is scored when wound healing of a full thickness epithelium covers the tail stump, preventing any regeneration. Note that the continued vacuolation of distal notochord cells produces a bulge under the wound epithelium, but no new cells are made. Arrowheads indicate the plane of amputation. **B)** Schematic of the two Tlr2 proteins coded by the L and S homeologues in *Xenopus laevis*. N, N-terminus; C, C-terminus, signal sequence and transmembrane spanning domains predicted by DeepTMHMM (Hallgren *et al*., 2022). Leucine rich repeats (LRR) and Toll-interleukin resistance domains (TIR) were mapped using NCBI conserved domains. sgRNA targeting the *Tlr2* genes are shown (rnk30 black arrows, rnk29 blue arrows) as approximate locations on the protein with * indicating the position of stop codons induced by frameshift Indels for each guide (185 for rnk30 in black, 536 for rnk29 in blue). CRISPants would be expected to have reduced Tlr2. **C)** Tail regeneration outcomes of stage 46 *Tlr2* CRISPant tadpoles from two sibships. Estimated editing percentage is averaged from three arbitrarily selected sibling embryos, calculated from Sanger sequence traces using TIDE. Ordinal χ^2^ was used to indicate significant differences in regeneration within each sibship, adjusted *p* values * *p.adj* <0.05, ** *p.adj* <0.01, **** *p.adj* <0.0001. Left sibship, N = 31 controls, 22 sgRNA rnk29 and 26 sgRNA rnk30 CRISPants. Right sibship, N= 36 controls, 40 rnk29 and 30 rnk30 CRISPants. **D)** Tadpoles from a third sibship were raised with or without 50 μg/mL gentamicin and tails amputated at stage 46, N=20 to 26 per group. Tadpoles were immediately either returned to media, or treated for two hours with 10 μg/mL *S. aureus* peptidoglycan and/or 10 μg/mL *E. coli* lipopolysaccharide before returning to media. They were scored for regeneration using categories in panel A. Ordinal χ^2^ was used to detect significant differences in regeneration within each sibship, columns that share a letter generated a *p.adj* >0.05. Raw data and analysis can be found in Tables S1-S5.

We have previously demonstrated the negative regeneration effects associated with use of gentamicin, an aminoglycoside antibiotic primarily active on Gram-negative bacteria and commonly used in rearing *Xenopus* tadpoles (Bishop and Beck, 2021; Chapman et al., 2022). We also showed that LPS, an important component of the Gram-negative bacterial outer membrane, can ‘rescue’ regeneration outcomes in tadpoles raised in gentamicin, as well as boost regeneration in tadpoles raised without antibiotics (Bishop and Beck, 2021; Chapman et al., 2022). LPS acts as a PAMP, binding to Tlr4 to activate the NF-κB transcription factor and downstream pro- and anti-inflammatory responses, depending on context (Wang et al., 2022). To see if providing other PAMPS at the time of tail amputation could also push more tails to regenerate in the refractory period, we added commercially purified peptidoglycan to regenerating tails, with or without LPS. Peptidoglycan and LPS in combination significantly increased the number of tadpoles undergoing tail regeneration in gentamicin raised tadpoles (Figure 1D).

### Can the skin microbiome predict tail regeneration outcomes?

As part of a previous study investigating how tadpoles obtain their skin microbiomes, we generated a large number of tadpoles from 16 sibships (Chapman et al., 2024). Embryos were raised without antibiotics and cultured in low density until stage 46. For each sibship, we removed the posterior third of the tail with a sterile scalpel blade, keeping the tail tip for 16S rRNA amplicon sequencing and recording the regenerative outcome for the corresponding tadpole after 1 week. Two 24 well plates were used to individually house tadpoles for each sibship while they underwent regeneration, and the first three of each plate were used for 16S rRNA amplicon sequencing in our previous study. The remaining 21 x 2 tadpoles (42) for sibships B4, B5, B7, B9, C1, C3, C4, C5, D1, D2, D3 and D4 (Figure 2A) were sequenced in a separate run and used in the present study. One individual in the C1 sibship was unable to be scored due to mortality and was excluded from the analysis. Ordinal χ^2^ tests showed that regenerative outcomes of plate replicates for each of the sibships were not different from one another, (apart from B4.1 vs. B4.2, *p*=0.04) and so data for both plates was pooled for analysis (Supporting Information S1). As expected, each sibship had a distinct regenerative profile (Figure 2A), with sibship regeneration rates - the percentage of tadpoles that made any attempt at regeneration (categories PB, PG and FR) of 40% - 100%, although all but two were more than 80%. Both sibships with notably poor regeneration rates were produced by females from Tank B, while sibships from Tank D mothers were consistently good regenerators, with regeneration rates of minimum 98%. Mothers within each tank were not necessarily related; however, mothers from Tank B were a significantly older cohort (13 years) than those in Tanks C and D (2-4 years).

**Figure 2:**
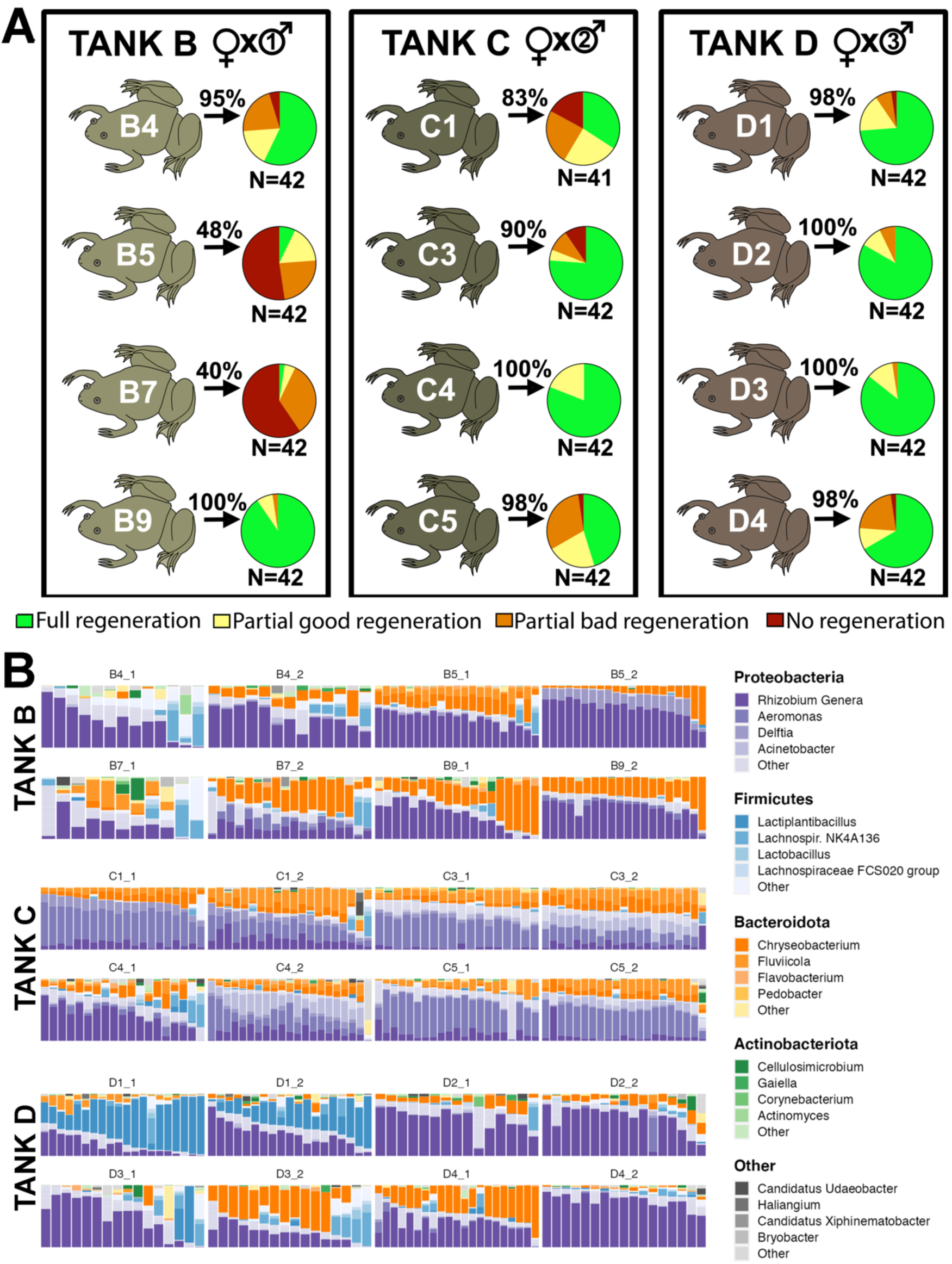
Schematic of experiment and tadpole regenerative outcomes. **A)** Four females from three aquarium tanks were induced to spawn and eggs fertilised with sperm from three males as indicated. Percentage of tadpoles that regenerated (FR, PG, PB) following tail amputation at stage 46 is indicated on the black arrows. Pie charts show how each tadpole cohort regenerated, and colour key shows examples of each category of regeneration after 5 days. N=42 tadpoles except C1 where one tadpole died and could not be scored. **B)** Tadpole tail 16S rRNA relative abundance of the four most abundant phyla, with the four most abundant Genera in each. Read counts were rarified to 1750 reads, with 438 samples retained for analysis. Raw data (2A) and summary data showing top genera for each tank (2B) are provided in Tables S6 and S7.

16S rRNA amplicon sequencing of 503 tail samples across the twelve sibships generated a mean of 11869 (+/- 10526, range 82 – 91228) reads per sample, with 438 samples passing rarefaction of 1750 reads (Figure S2). Four bacterial phyla dominated the tadpole tail skin: Proteobacteria were most abundant, followed by Bacteriodota, Firmicutes and Actinobacteriota (Figure 2B).

We next looked for associations between microbial genera and regeneration outcomes based on the 16S data. Regardless of regeneration outcome, tadpoles were dominated by *Rhizobium* taxa, with *Chryseobacterium*, *Aeromonas* and *Fluviicola* also being common across all outcomes (Figure 3A). We tested for correlations between microbial community composition and regeneration success (Figure 3B). Plates were clustered based on Spearman’s correlation coefficient, calculated on mean relative abundance of microbial genera. *Rhizobium* and *Chryseobacterium* spp. were generally most abundant in tadpoles with mothers from Tanks B and D, with *Aeromonas* being dominant in most Tank C samples.

**Figure 3.**
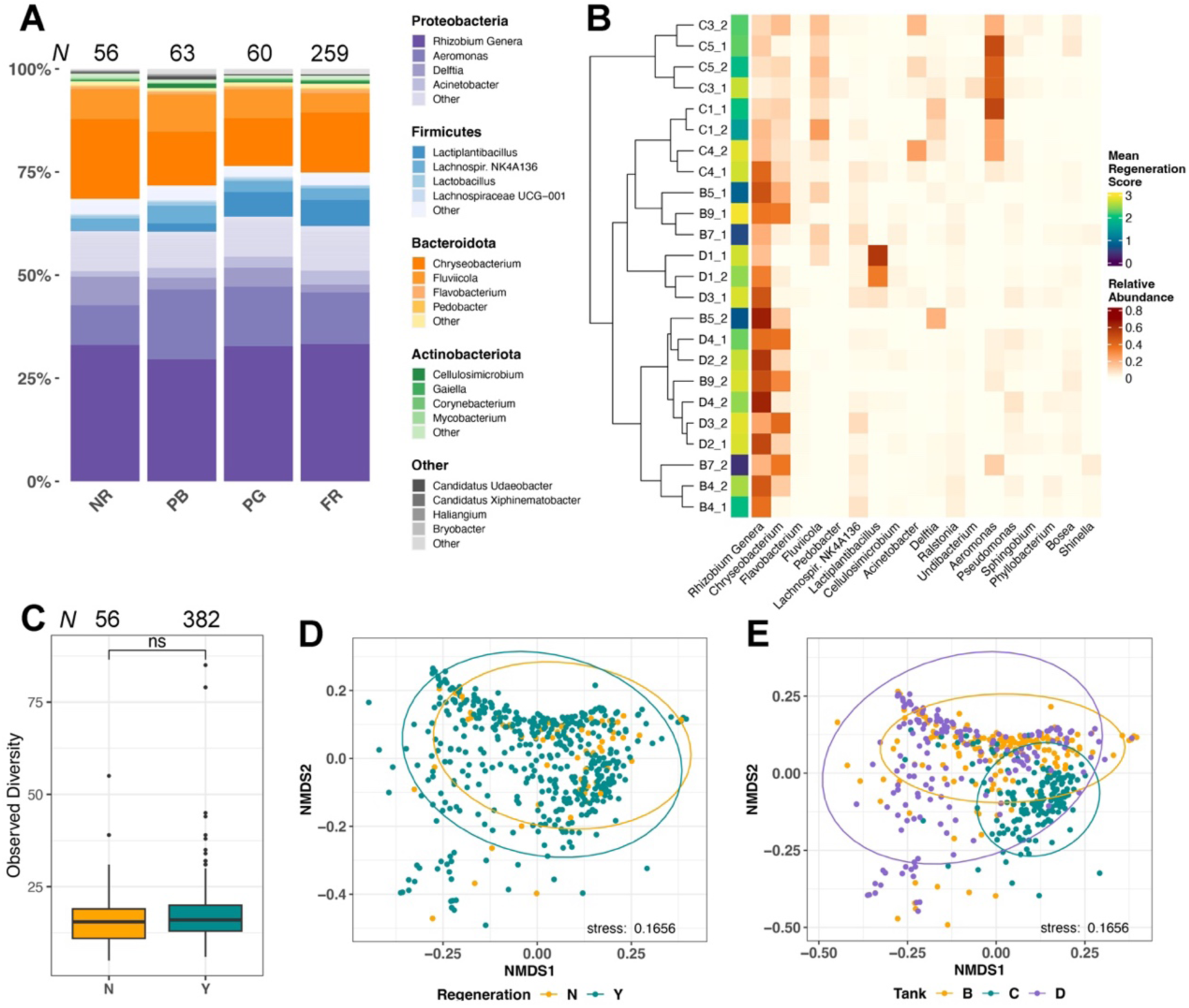
The composition of skin microbiomes in tadpoles with different regeneration outcomes. **A.** Relative abundance (rarefied at 1750 reads) of bacterial taxa based on regeneration score. NR = No Regeneration, PB = Partial Bad Regeneration, PG = Partial Good Regeneration, FR = Full Regeneration. Sample size *N* is shown at the top of each bar. **B.** Heatmap showing mean relative abundance of microbial genera in tadpole sibships. Sibships were divided into two plates each. Taxa shown are those that comprise a minimum 2.5% of at least one sample. Tadpole plates are depicted on the y-axis and are clustered according to Spearman’s correlation coefficient calculated on microbial mean relative abundances. Regeneration scores were assigned for each tadpole (0 = no regeneration, 1 = partial bad regeneration, 2 = partial good regeneration, 3 = full regeneration) and a mean score calculated for each plate**. C)** Observed alpha diversity of NR non-regenerating (defined as N) samples was not different from regenerating samples (PG+PB+FR, defined as Y) (*p*<0.05 = ns, Wilcoxon rank test) Sample size *N* is shown at the top for each bar. **D, E)** Non-metric multidimensional scaling plots calculated on weighted UniFrac distances, with stress 0.1656. **D)** Non regenerators (N) vs. all regenerators (Y) **E)** Offspring of females in tanks B and D group more closely than those in tank C. Data for A is in Table S8. Statistical analysis of C, D and E can be found in Tables S9-S11.

Alpha diversity metrices (Observed, Shannon, Faith’s Phylogenetic) showed no significant differences based on regeneration outcome (Observed, Figure 3C, Shannon and Faiths Figure S3). NMDS (beta diversity), based on weighted UniFrac distances, did not show clear definition of regeneration groups (Figure 3D), although a Hopkins statistic of 0.91 indicates some clustering and silhouette analysis supports two clusters in the data (Figure S4). This is most easily explained as tank-of-origin effect as the tail microbiomes of tadpoles from mothers in C clustered separately from B and D (Figure 3E). PERMANOVA indicated a significant effect of regeneration (although this accounted for less than 1% of weighted UniFrac distance variation seen; R^2^ = 0.006, p = 0.001) and sibship/mother ID (35.2%; R^2^ = 0.352, p = 0.001), as well as a significant interaction between mother and plate ID (15.2%; R^2^ = 0.152, p = 0.001).

### Regeneration outcome does not depend on a specific taxon, but several genera were associated with regenerative success

To test whether microbial genera were consistent predictors of regeneration outcome, we built a random forest classification model. The model compared samples based on any (FR, PG, PB categories) vs. no regeneration (NR category), and achieved a mean receiver operating characteristic area under curve on the test dataset of 0.68, compared with 0.67 on the training dataset, indicating minimal overfitting. The 20 best predictors of classification, based on mean decrease in accuracy, are shown in Figure 4A. *Delftia* had the greatest effect on model accuracy, however this was low at approximately 0.08%. Correlation analysis/differential abundance analysis of the tail 16S rRNA data with either ANCOMBC2 or MaAsLin2 analysis identified four genera (*Lactiplantibacillus*, *Pseudomonas, Sphingobium, Cupriavidus*) as being significantly more abundant in tail samples from tadpoles that went on to regenerate (PB, PG, FR categories) vs. non regenerators (NR) (Table 1). Two genera were found to be significantly lower in successful regenerators *(Klebsiella, Delftia*). However, when the identity of the mother/sibship was added as a fixed effect alongside regeneration outcome, these genera (with the exception of *Klebsiella,* Table S13) were no longer differentially abundant between regeneration categories.

**Figure 4.**
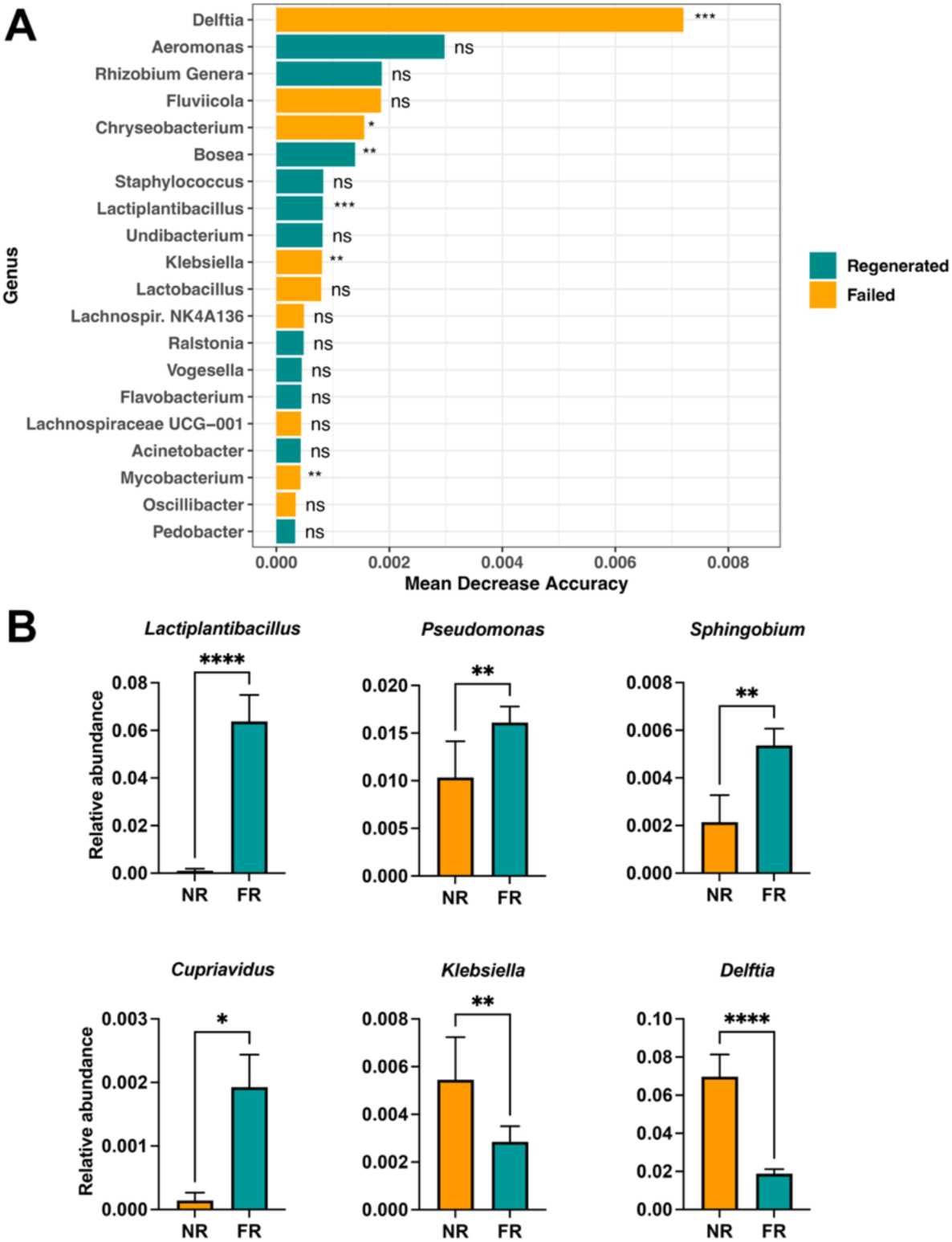
*Delftia* species are the highest predictors of regeneration, but several other genera are also more or less abundant in non-regenerators. **A).** Random Forest analysis. Top 20 most important taxa for classifying tadpoles into regenerator or non-regenerator categories, based on mean decrease in accuracy in random forest analysis. Taxa have been split into two groups for visualisation; those in the “Failed” category (tadpoles scoring as NR) had higher mean abundance in non-regenerating tadpoles, while those in the regenerated group (all tadpoles scoring PB, PG or FR) were more abundant in regenerators. Wilcoxon rank sum test was used to identify significant difference between regenerating and non-regenerating groups. **B)**. Bar plots (mean, sem) of relative abundance of the six genera found to be differentially abundant between NR (*N*=56 samples) and FR (*N*=259 samples) groups using MaAsLin2 or ANCOMB2. Mann-Whitney U was used to test for significant differences between the NR and FR groups,* P<0.05, **p<0.01, ***p<0.001 **** p<0.0001. Raw data and statistics can be found in Tables S12 and S14-19.

**Table 1.** Differential abundance of microbial taxa found on regenerating (PB, PG, FR) vs. non-regenerating (NR) tadpole tails.

| Genus | ANCOMBC2 |  |  | MaAsLin2 |  |  |
| --- | --- | --- | --- | --- | --- | --- |
|  | p val | q val | Higher in: | p val | q val | Higher in: |
| *Lactiplantibacillus* | 0.000 | 0.010 | Regeneration | 0.000 | 0.002 | Regeneration |
| *Pseudomonas* | 0.019 | 0.205 | Regeneration | 0.003 | 0.042 | Regeneration |
| *Sphingobium* | 0.026 | 0.236 | Regeneration | 0.000 | 0.042 | Regeneration |
| *Cupriavidus* | 0.012 | 0.162 | Regeneration | 0.005 | 0.051 | Regeneration |
| *Klebsiella* | 0.012 | 0.162 | Non-regeneration | 0.009 | 0.085 | Non-regeneration |
| *Delftia* | 0.000 | 0.006 | Non-regeneration | 0.000 | 0.002 | Non-regeneration |

\* cutoff FDR $q < 0.25$ , N=503 samples

As non-regenerators (NR) account for only 63 of the 503 samples, so we also compared relative abundance of group to the full regeneration (FR) group (295 samples). MaAsLin2 analysis of NR vs FR identified twelve differently abundant genera, and ANCOMBC2 also identified a subset of seven of these. These core differentially abundant genera matched those identified when NR were compared to all regenerators, as above, with the addition of *Vogesella* to the “more abundant in regenerators” group (Table 2). However, after the additional of mother identity as fixed effect, none of these directional effects were preserved (Table S13).

**Table 2.** Differential abundance of microbial taxa found on full regenerating (FR) vs. non-regenerating (NR) tadpole tails.

| Genus | ANCOMBC2 |  |  | MaAsLin2 |  |  |
| --- | --- | --- | --- | --- | --- | --- |
|  | p val | q val | Higher in: | p val | q val | Higher in: |
| *Lactiplantibacillus* | 0.000 | 0.005 | FR | 0.000 | 0.001 | FR |
| *Pseudomonas* | 0.005 | 0.097 | FR | 0.001 | 0.009 | FR |
| *Sphingobium* | 0.008 | 0.110 | FR | 0.001 | 0.009 | FR |
| *Acinetobacter* | ns | ns | - | 0.021 | 0.143 | FR |
| *Bosea* | ns | ns | - | 0.012 | 0.131 | FR |
| *Vogesella* | 0.025 | 0.226 | FR | 0.001 | 0.111 | FR |
| *Ralstonia* | ns | ns | - | 0.038 | 0.207 | FR |
| *Cupriavidus* | 0.016 | 0.177 | FR | 0.007 | 0.082 | FR |
| *Shinella* | ns | ns | - | 0.043 | 0.213 | NR |
| *Thermoanaerobacterium* | ns | ns | - | 0.051 | 0.236 | NR |
| *Klebsiella* | 0.031 | 0.243 | NR | 0.027 | 0.163 | NR |
| *Delftia* | 0.000 | 0.000 | NR | 0.000 | 0.000 | NR |

\* cutoff FDR $q < 0.25$ , N= 358 samples

The relative abundance of the seven genera identified by the differential abundance analyses are compared for NR vs. FR in Figure 4B. The detected relative abundance of 16S rRNA mapping to *Lactiplantibacillus*, a Gram-positive genus, was an average of 61 x higher on tail samples from tadpoles that went on to regenerate a full tail (FR) than in the NR group. It should be noted though that *Lactiplantibacillus* was in high abundance in the D1 sibship (Figure 2B). FR samples also had a 11.6 x higher mean relative abundance of reads mapping to *Cupriavidus*, 2.5 x more *Sphingobium*, and 1.6 x more *Pseudomonas.* In contrast, NR tail samples had 3.7 x higher mean RA of *Delftia* and 1.9 x more *Klebsiella* (Figure 4B).

### Tadpole tail regeneration in refractory stages does not correlate with maternal relatedness

Maternal lineage was the biggest correlate with regenerative outcome, but it is not clear whether this is due to genetics or to transfer of microbiomes described in Chapmen et al (2024). To see how closely related our sibships were, we sequenced the mitochondrial genomes of the twelve females in the study. These were used to construct a Maximum Likelihood tree (Figure 5). These genomes were aligned to the *X. laevis* reference MN259072.1 (Evans et al., 2019). Unsurprisingly, since our colony has been closed for 20 years, mothers formed two major clades, with one individual (B7) forming a clade of its own. There was no obvious correlation between mean regeneration score of the tadpoles and the phylogenetic position of their mothers. This is illustrated by the B9, D3, and C4 sibships, all of which underwent regeneration, having an identical mitochondrial genome to the poorly regenerating B5 sibship.

**Figure 5.**
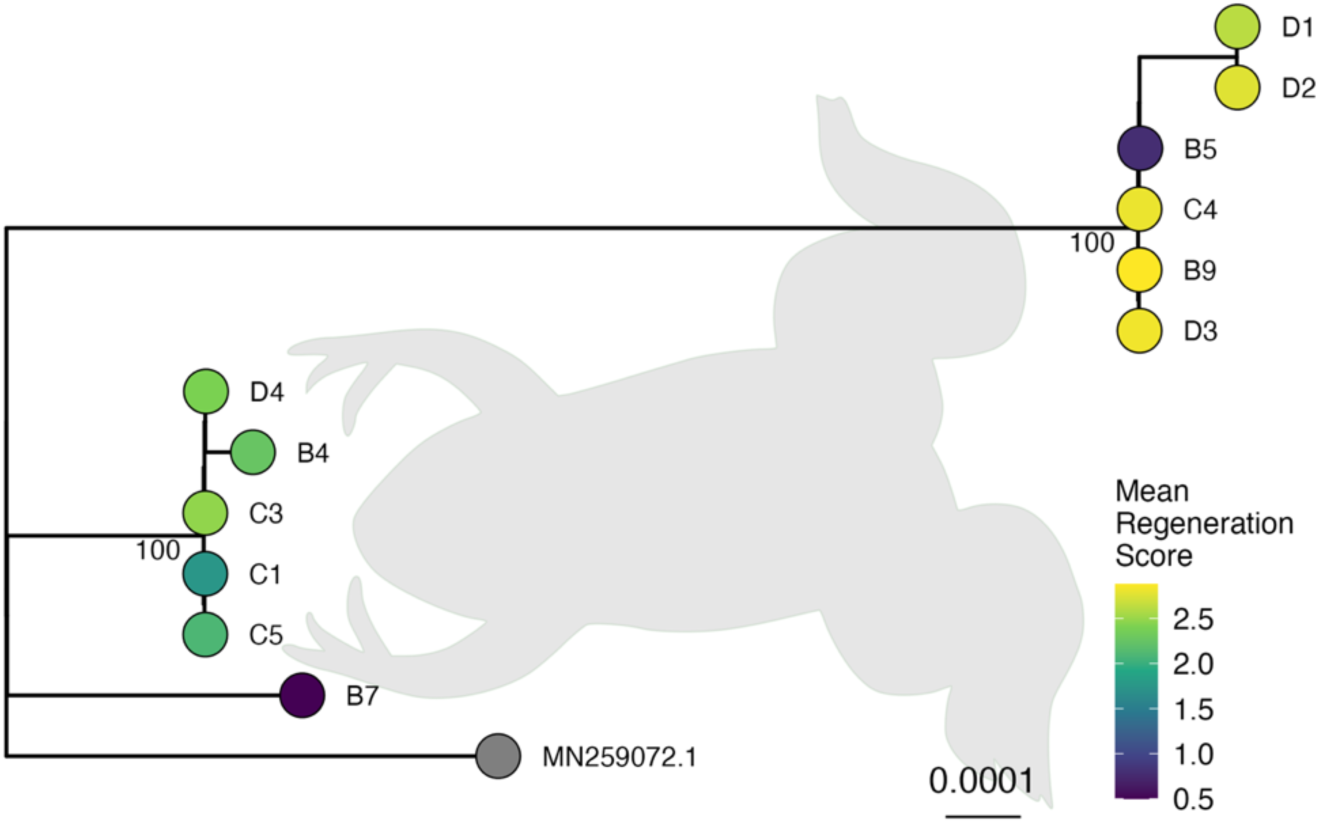
Maximum likelihood tree inferred from the mitochondrial genome. Scale bar indicates the mean number of substitutions per base. Regeneration scores were assigned for each tadpole (0 = no regeneration, 1 = partial bad regeneration, 2 = partial good regeneration, 3 = full regeneration) and a mean score calculated for each mother/sibship. Reference genome MN259072.1 (Evans *et al*., 2019) was used as an outgroup. Sequences are available at GenBank, accession numbers PP467366-PP467377.

### Bacterial skin load is unlikely to influence regenerative outcomes

We had previously proposed a role for Gram-negative bacteria on the skin providing a source of LPS which could trigger regenerative responses via interaction with the innate immune signalling pathway and TLRs. LPS triggers the same immune responses whether the bacteria are dead or alive, so to be effective, antibiotics should be added to the embryo media as early as possible. We used a commercial endotoxin (LPS) detection kit to directly quantify the amount of endogenous LPS associated with a single tadpole tail tip (Figure 6). Tail tips from tadpoles raised without antibiotics varied most in the level of LPS/endotoxin, ranging from 0.08 to 3.14 EU/mL. In contrast, LPS levels from tadpoles from the same sibships that had been raised in gentamicin were consistently below 0.5 EU/mL. However, there was no difference in the mean LPS levels of tail tips from tadpoles that regenerated (FR) compared to those that did not (NR) for either control or gentamicin raised tadpoles (Figure 6A).

**Figure 6.**
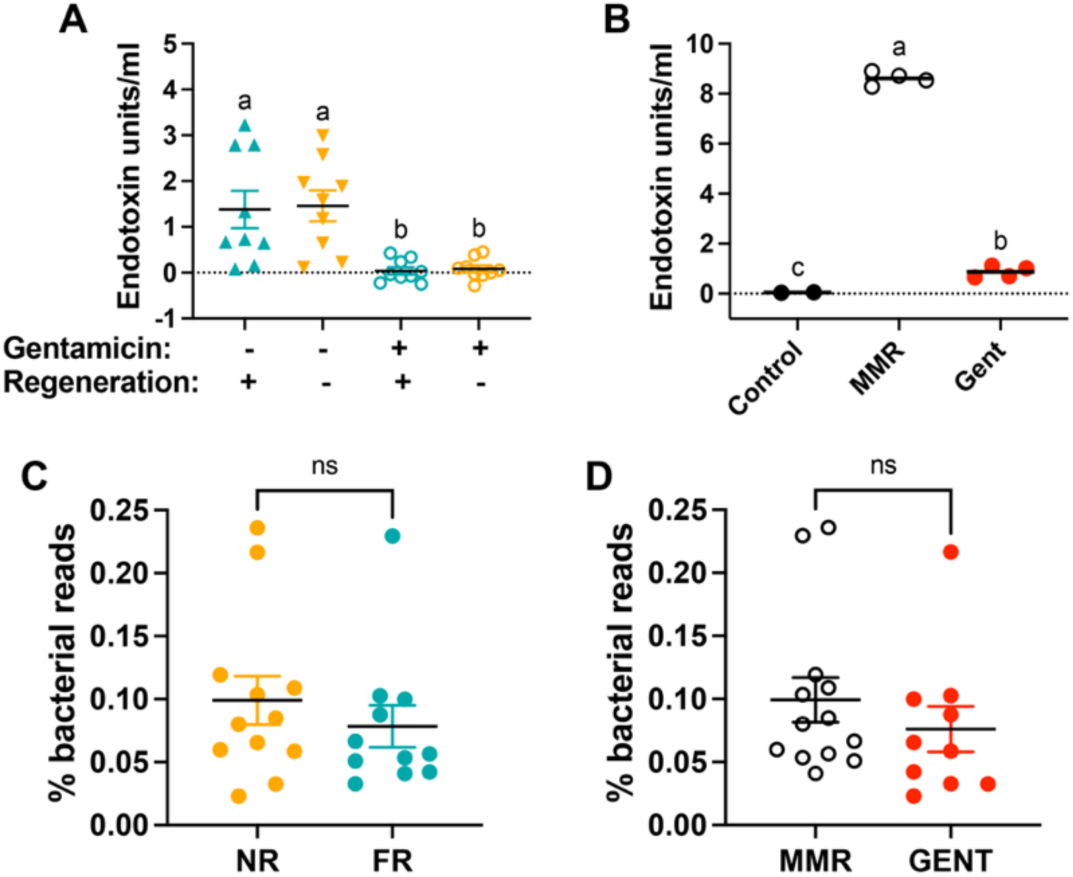
Bacterial skin loading does not predict regenerative outcomes. A, **B)** Direct measurement of LPS (endotoxin), shown as scatterplots. Datapoints are average of three technical assay replicates and each point is a biological replicate. Groups were compared using 1-way ANOVA with Tukey’s post hoc testing of all means. Groups that do not share a letter (compact letter display) are significantly different from one another (*pAdj* <0.05). **A)** Endotoxin measurements from individual tadpole tail samples, raised with or without gentamicin, that went on to fully regenerate (FR) or did not regenerate (NR). N=8 tadpole tails per group. **B)** Endotoxin from unused MMR media (N=2) compared to media in which tadpoles were raised with (N=4) or without gentamicin (N=4). **C, D)** Scatterplots showing the percentage of reads mapping to bacterial sequences of shotgun genome sequenced tail tip DNA **C)** Tail tips from tadpoles that did not regenerate (NR, N=12) compared to those with full regeneration (FR N=11), Mann-Whitney test, ns= non-significant. **D)** Tail tips from tadpoles raised with (N=10) or without (N=13) the antibiotic gentamicin. Mann-Whitney test, ns= non-significant. Raw data and statistics can be found in Tables S20-S26.

We therefore saw no evidence that LPS levels associated with tadpole tail skin could predict or influence regeneration of an individual tadpole. However, when we measured levels of LPS from the tadpole’s media, a different story emerged (Figure 6B). Much higher levels of LPS were detected in tadpole MMR media when gentamicin had not used (mean 8.61 +/- 0.13 EU/mL, *N*=4 plates). This was ten-fold higher than was detected in media from plates where gentamicin had been used to grow the same batches of tadpoles (mean 0.87 +/- 0.11 EU/mL, N=4 plates). No LPS was detected in our tadpole culture medium (0.1 x MMR, mean <0.05 EU/mL).

We next attempted to semi-quantify bacterial load using shotgun short read sequencing of 24 tail samples from sibships B5, C1 and D1. UMI (unique molecular indicator) barcodes were used to control for PCR amplification bias, and C1 samples were also spiked before extraction using a known number of three “foreign” bacterial species. As expected, most of the reads were from the host, with an average of 0.12% reads mapping to bacteria (mean 72,740 bacterial reads per sample). The proportion of total reads from each library that mapped to bacteria did not differ between tail samples from tadpoles that subsequently underwent full regeneration or no regeneration (Figure 6C, mean NR 0.10 +/- 0.02 vs. FR 0.08 +/- 0.02%). One NR sample from B5 had much higher reads than any others and was removed from analysis as an outlier; this did not alter the conclusions. Taken together with the LPS measurement data, this supports the idea that tadpole tail regeneration is not activated by a threshold number of bacteria on the skin.

The shotgun sample set also included tail tips from gentamicin-raised tadpoles from sibships D1 and C1, which we predicted would have a lower bacterial read count than their naturally raised siblings. However, there was no difference in the percentage of bacterial reads from gentamicin raised tail tips vs. controls (Figure 6D, control mean (0.10 +/- 0.02%, N=13) and gentamicin mean (0.08 +/- 0.02%, N=10). To be sure that this result was indicative of bacterial loading, we also analysed the C1 samples using standard “spike-in” non commensal species reads to quantify bacterial DNA, and found no differences between NR or FR samples from tadpoles raised either with or without gentamicin (Figure S5).

### Regenerative success is associated with an increased proportion of Gram-positive commensal bacteria on tadpole tail skin

We were unable to identify a signature microbiome or a single taxon associated with successful tadpole tail regeneration. However, noting the positive effect of adding exogenous PAMPs LPS and PG, we wondered if the relative abundance of Gramnegative taxa (which contribute LPS) to all Gram-positive taxa (which contribute the most peptidoglycan) might influence regenerative outcomes. We therefore calculated the relative abundance of total bacterial reads comprised by Gram-positive genera in the mapped 16S tadpole tail sample set. When these were grouped by regeneration outcome (NR, PB, PG or FR), the proportion of Gram-positive reads was only significantly higher in tail tips from tadpoles that went on to undergo full regeneration (FR vs. NR, Wilcoxon’s Rank Sum, with continuity correction *p* = 0.012; Figure 7).

**Figure 7.**
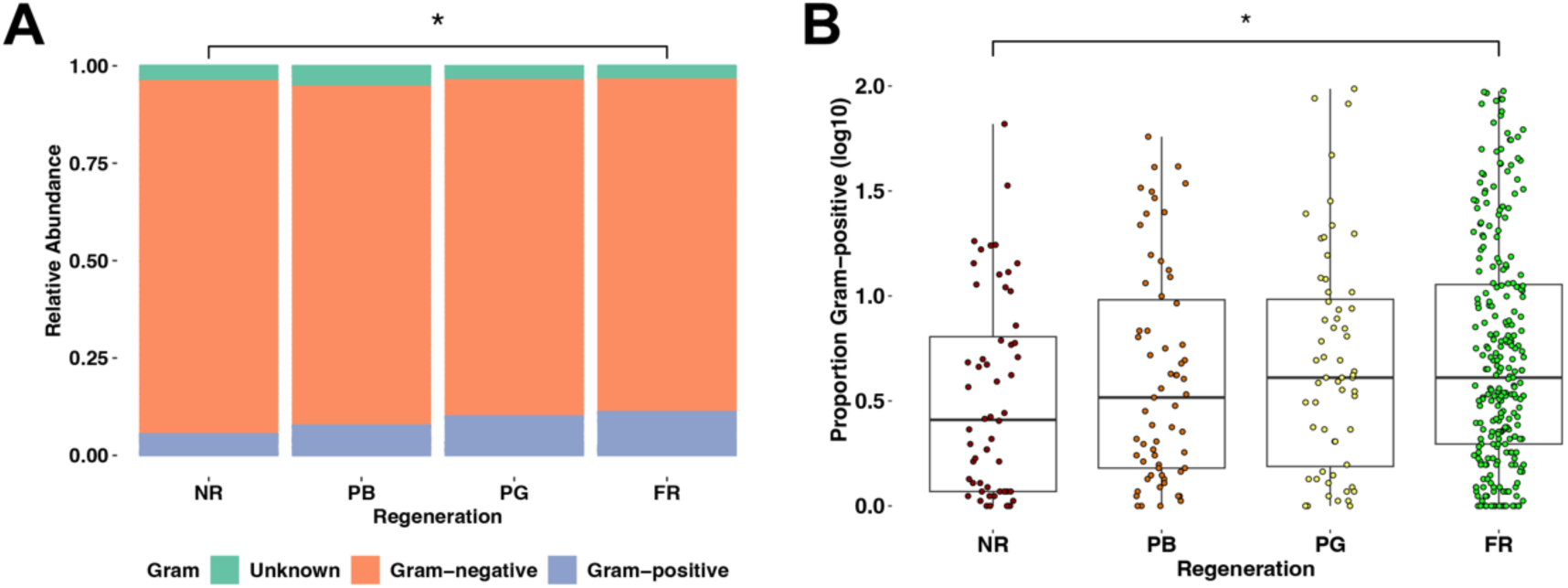
Regeneration is associated with increased proportion of Gram-positive bacteria in skin microbiome. **A)** Relative abundance plot of Gram-positive, Gram-negative or unknown (could not be assigned to either group) 16S rRNA reads, from all 438 samples from the 12 sibships. **B)** Proportion of Gram-positive bacteria (log10 transformed) on tails from tadpoles that did and did not attempt regeneration after amputation. A significant difference was observed based on Wilcoxon’s Rank Sum (p = 0.012). Sample sizes: *N*= 56 (NR), 63 (PB), 60 (PG) 259 (FR). Raw data and analysis can be found in Supporting Information file S1.

Due to this unexpected finding, we investigated the effect of adding an antibiotic that was selective for Gram-positive inhibition. Vancomycin targets actively dividing Gram-positive bacteria, preventing cell wall cross links from forming (Jordan, 1961). Four sibships of tadpoles, unrelated to the microbiome study animals, were raised either with no antibiotic, or with 50 μg/mL gentamicin or vancomycin added to the media from the 4-cell stage. Successful tail regeneration at stage 46 was significantly less likely in tadpoles raised in either antibiotic across all four sibships (Figure 8).

**Figure 8.**
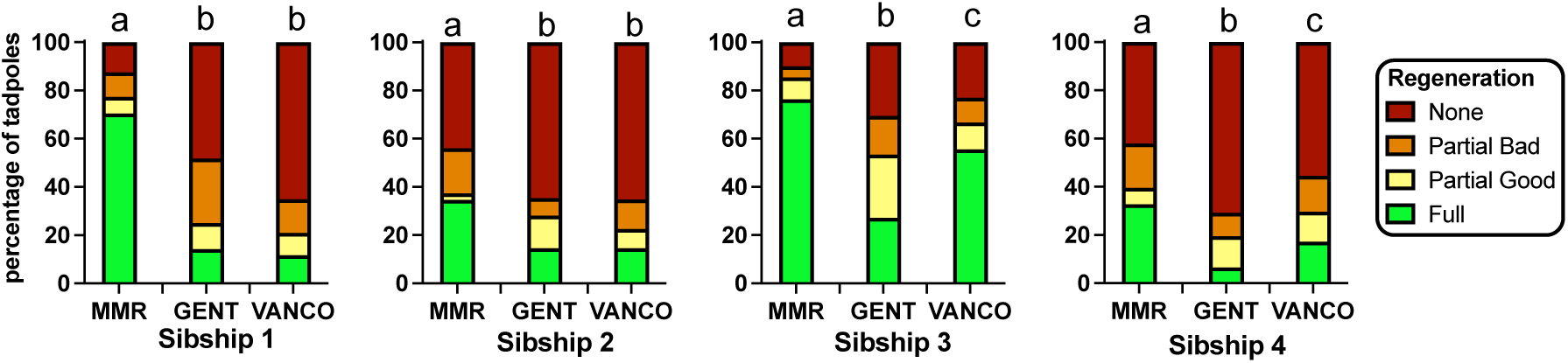
Raising tadpoles in the presence of the Gram-positive selective antibiotic vancomycin reduces regeneration success. Four tadpole sibships were raised without any antibiotic (MMR) or in MMR containing 50 μg/mL gentamicin (GENT) or vancomycin (VANCO). Regeneration assays were conducted and scored as in Figure 1. Sibships 1 and 2 were *N*=56-89 for each treatment, data combined from 3 plates, and sibships 3 and 4 were *N*=110-128 tadpoles per treatments, data combined from 4 plates. Ordinal χ^2^ was used to detect significant differences in regeneration within each sibship, columns that share a letter generated a *P adj* >0.05 and are not significantly different. Raw data and analysis can be found in Tables S28 and S29.

## Discussion

### Activation of the innate immune system by PAMPS promotes regenerative outcomes

In the past three decades, it has become increasingly clear that metazoans and their microbial commensals have evolved together, and that disruption of this “holobiont” can lead to chronic disease states, often linked to inflammation (Postler and Ghosh, 2017; Prescott et al., 2017). In humans, these inflammatory disorders can be primed by altered signalling between the microbiome and host, either during the development of the immune system or later in life. We have previously shown that growing tadpoles in antibiotics like gentamicin, which is common practice in *Xenopus* labs, both reduces the number of tadpoles that undergo regeneration and modifies the composition of the skin microbiome (Bishop and Beck, 2021; Chapman et al., 2022). Adding back heat-killed bacteria, commercially prepared LPS (Bishop and Beck, 2021), LPS from commensal species (Chapman et al., 2022), or commercially prepared peptidoglycan (this study) at the time of tail amputation counters the negative effect of raising tadpoles in antibiotics. These results support a model whereby the refractory period, where *X. laevis* tadpole tails often undergo full thickness skin wound healing rather than regeneration, is the result of interfering with normal bacterial growth. This effect can be rescued in the presence of the innate immune system-activating PAMPs LPS or PG in the tadpole’s media. PAMPs are small molecules found in microbes but not in the host, which can bind to receptors of the innate immune system, forming a first line of defence against infection. We propose that when the tadpole tail is amputated, the barrier normally provided by the skin is breached, allowing “non-self” PAMPs to interact with the host’s innate immune pattern recognition receptors (PRRs) on newly exposed tissue-resident myeloid (“professional”) phagocytes, triggering an altered inflammatory response.

Vertebrate PRRs comprise five different protein homology groupings, including Toll-like receptors reviewed in (Li and Wu, 2021). We hypothesised that Tlr4, Tlr5 and Tlr2 were most likely to be involved with sensing microbial PAMPs since these three receptors are presented on the surface of myeloid phagocytes. We previously showed a reduced regenerative response in tadpoles with CRISPR/Cas9 knockdown of *tlr4.S* (Chapman et al., 2022). Tlr4 homodimers are the main PRR for LPS, and *X. laevis* has only one homeologue of this receptor, *tlr4.L*. Tlr5 specifically binds flagellin, a component of bacterial flagella (Hayashi et al., 2001), and is encoded by the *X. laevis tlr5.L* gene. Here, we looked at the effect of knocking down Tlr2, which has both an L and S homeologue in *X. laevis*. Edits causing premature truncation of the Tlr2 protein about a third of the way down the extracellular, ligand binding N-terminal leucine rich repeat domain led to almost none of the crispant tadpoles regenerating a tail. Tlr2 however, does not commonly act as a homodimer, instead partnering with TLR6 to bind bacterial peptidoglycan from Gram-positive pathogens (Ozinsky et al., 2000). We interrogated single cell RNA sequencing data of *X. laevis* tail regeneration (Aztekin et al., 2019) and found that *tlr2.L, tlr2.S, tlr4.S, tlr5.L* and *tlr6.L* were all expressed in “myeloid1” cells associated with inflammatory responses. We suggest that knocking down *tlr2* abrogates the ability of these cells to sense peptidoglycan from Gram-positive commensals, since Tlr2/6 heterodimers will be reduced. While we cannot rule out the possibility that Tlr2 homodimers, which bind other PAMPS, cause the loss of regenerative response, others have shown that only the heterodimers activate the pro-inflammatory transcription factor NF-κB (Ozinsky et al., 2000).

### Gram-positive bacteria may influence regeneration

Our study of the bacterial commensals of 503 individual tadpole tails found no single bacterial genus to be predictive of regenerative success. A common microbiome comprised of Gram-negative *Rhizobium, Chryseobacterium*, *Aeromonas* and *Fluviicola* genera was seen across all tadpole sibships, in agreement with previous results (Chapman et al., 2024). However, when looking at the cohort overall, 16S rRNA reads from the Gram-positive *Lactiplantibacillus* were detected at far higher relative abundance in tail samples from tadpoles that went on to regenerate well.

*Pseudomonas* species relative abundance was also associated with positive regenerative outcomes both here and in our previous study with gentamicin raised tadpoles of the same stage (Chapman et al., 2022). Two other genera were significantly more abundant on tail tips from untreated tadpoles that subsequently underwent regeneration: *Sphingobium* and *Cupriavidus*. Conversely, *Delftia* and *Klebsiella* species were associated with poor regenerative outcomes in both random forest and relative abundance analyses. *Delftia* was the strongest predictor of regenerative outcome in the random forest model, but the effect of removing it was still <0.1%.

Our current findings do not support a role for Gram-negative derived LPS levels in determining the regenerative status of an individual tadpole. The number of Gram-negative bacteria colonising the skin, inferred from either normalised sequence reads, spike-in quantification, or direct LPS detection, does not predict regenerative success. Neither did we find any evidence that raising in antibiotics reduced bacterial read number, even though it did reduce the amount of LPS detected on tadpole tails. This suggests that there are limited opportunities for bacteria to colonise the tadpole tail skin, and when some species are selected against others move in. Competition for places is also initially bottlenecked as each mother will only pass on a subset of her bacterial commensals to tadpoles via the egg jelly (Chapman et al., 2024). From our measurements of endogenous LPS in tadpole tail and media, it is likely that the level of exposure to dead or dislodged bacteria is much higher in naturally raised tadpoles. This could result in higher levels of PAMP exposure on tail amputation which in turn raises the chances of regeneration in a batch of tadpoles swimming in shared media. The individual tadpole’s microbiome, which is mostly Gram-negatives, may therefore become less influential in naturally raised tadpoles.

Surprisingly, we found that relative abundance of Gram-positive bacterial reads correlated to better regenerative outcomes overall. The possibility that Gram-positive species can drive regeneration is supported by our observations that *tlr2* crispants almost never regenerate, and that peptidoglycan addition increased the chance of regeneration. When we agglomerated all 16S rRNA reads by Gram-positive, Gram-negative or unknown status, a trend of increasing Gram-positives with better regeneration and a significant difference between the non-regenerators (NR) and full regenerators (FR) was revealed. To see if this was biologically relevant, we tested a Gram-positive specific antibiotic, vancomycin, which targets cell wall synthesis, and found that this had a consistent effect on inhibiting regeneration. Vancomycin is too large to pass through the aqueous porin channels of Gram negative bacteria and is therefore only supress proliferation of Gram-positives (Masi et al., 2019). Despite this limitation, tadpoles raised in vancomycin regenerated at similarly low rates to those raised in the broad-spectrum aminoglycoside gentamicin. Taken together, our findings suggest a model whereby refractory stage tadpoles can regenerate their tails at much higher rates under natural conditions which enable the establishment of a microbiome via vertical maternal transfer.

### Conclusions and Study limitations

We have sequenced bacterial 16S rRNA from a large cohort (n=503) of tadpole tail samples of known parentage and mapped these to the regenerative outcome of each tadpole. While we were unable to find a typical microbiome associated with regeneration, we reveal that regenerative success correlates with increased proportions of reads from Gram-positive bacteria, which have higher levels of the PAMP peptidoglycan. This PAMP is detected by Tlr2/Tlr6 heterodimers on professional phagocytes (Ozinsky et al., 2000). These myeloid calls have been shown by others to be critical for tadpole tail regeneration (Aztekin et al., 2020; Pentagna et al., 2021). Tlr2 CRISPR knockdown tadpoles are less able to regenerate tails, and adding peptidoglycan increases regenerative success. While we have not directly demonstrated the activation of Toll-like receptors by endogenous PAMPs, the finding that Gram-positive specific antibiotic vancomycin also prevents regeneration also supports our model that bacterial PAMPs detected by exposed myeloid cells modulate inflammatory responses, leading to different regenerative outcomes. However, mice born from mothers treated with antibiotics during pregnancy have altered innate immune systems due to effects on myeloid cell development (reviewed in Li and Wu (2021)). Therefore, antibiotic raised tadpoles may have fewer tissue resident myeloid phagocytes to respond to PAMPs, and we cannot rule out this possibility.

## Materials and Methods

### Animal ethics

*Xenopus laevis* adults were held under University of Otago Animal Ethics Committee animal use procedures AUP22-24, and eggs and embryos were produced according to AUP19-01 and AUP22-12.

### Animal husbandry, egg collection, and fertilisation

The *Xenopus laevis* colony at University of Otago has been closed since 2004 and was established from adult frogs from the former colony at University of Bath, UK. All frogs were bred in-house from these founders. Animals are housed in recirculating aquaria supplied with carbon-filtered tap water as described in Chapman et al. (2022). Egg laying was induced by injecting 500 IU Chorulon per 75 g of adult female bodyweight into the dorsal lymph sac with a 27.5-gauge injection needle, 16 hours before required. Females were then moved to individual 1 L tanks containing 1 x MMR (Marc’s modified ringers: 10 mM NaCl, 0.1 mM MgSO_4_.6H_2_O, 0.2 mM CaCl_2_, 0.5 mM HEPES, 10 μM EDTA, pH 7.8) from which eggs were collected before *in vitro* fertilisation using fresh testes from a euthanised adult male *X. laevis.* Jelly coats were removed 20-30 minutes after fertilisation, by swirling gently in a 2% solution of L-Cysteine at pH 7.9 and rinsed 3 x with 1 x MMR. At the 4-8 cell stage embryos were transferred into 0.1 x MMR, with antibiotic treatment (if used) added at this stage. Embryos and tadpoles were staged according to Nieuwkoop and Faber’s staging series (Nieuwkoop and Faber, 1994) and raised in incubators at 18 to 23°C.

### Tail regeneration assays

To test the effects of Tlr2 ligands (peptidoglycan) on tail regeneration while accounting for sibship effects, we carried out tail regeneration assays as outlined below. Tail regeneration assays were carried out as previously described (Chapman et al., 2022) but procedures are summarised below.

#### Peptidoglycan challenge

Embryos from each sibship were split into two groups at the 4-cell stage with half being raised to NF stage 46 in the presence of 50 μg/ml of gentamicin sulphate. The medium was changed every 48 hours. Once the tadpoles had reached NF stage 46, they were briefly anaesthetized with 1/4000 MS222 (tricaine) to enable removal of the posterior one third of their tail using a sterile scalpel blade. Purified peptidoglycan (PG) from *Staphylococcus aureus* (Sigma 77140) and lipopolysaccharide (LPS) from *Escherichia coli* 055 : B5 (Sigma L2880) stocks were dissolved in endotoxin free water at 1 mg/mL. Within 2 minutes of tail amputation, tadpoles were moved in batches of 12 to sterile 5 mm petri dishes containing 10 ml of 0.1 x MMR and 10 μg/mL of LPS, PG, or both, for 2 hours. Untreated controls were incubated in the same way, but with no LPS or PG added. Following treatment, the tadpoles were returned to 10 cm petri dishes containing 30 mL of 0.1 x MMR and incubated at 18°C. After one week, the tadpoles were assessed for regeneration. Outcomes were scored as full regeneration (FR: all tail tissues reformed), partial good regeneration (PG: all tissue present, small defects such as partially missing ventral fin or tail bend), partial bad regeneration (PB: one or more tissues not formed), or no regeneration (NR: full thickness sin covers tail stump, no regenerated tissues). Examples of each category can be found in Figure 1A.

#### Replicate data for female frogs

Replicate spawning data was obtained for the offspring of 8 of the 12 female frogs in the study. Females were induced as previously described, in either 2021 or 2022, and eggs collected for fertilization. The male used was different in each case, so the resulting tadpoles are half-siblings to the 2020 cohort. Tadpoles were raised at 18°C in incubators in groups of 30 until tadpoles reached the refractory stage 46, at which point tail amputations were carried out. Regeneration was again scored after 1 week.

#### Antibiotic tests

Embryos from unrelated female frogs were used for this test. Four sibships were obtained and each was split into three treatment groups at the 4-cell stage. From this stage they were raised with no antibiotic, 50 μg/mL of gentamicin sulphate (Sigma G1914), or 50 μg/mL vancomycin (Merck: 1709007). Media was changed every 48 hours with fresh antibiotic where applicable, and tadpoles were assayed for tail regeneration at stage 46 as above, except that antibiotics were continued for 24 hours after tail amputation.

### CRISPR/Cas9 disruption of Tlr2

Unique sgRNA sequences targeting *Xenopus laevis Tlr2.S* and *L* homeologues were identified using ChopChop v2 (Labun et al., 2016), Table 3). InDelphi (Shen et al., 2018) was used to predict frameshifts resulting from editing (Table 3). EnGen sgRNA oligo designer v1.12.1 tool (NEB) was used to generate 55 bp oligos which were synthesized by Integrated DNA Technologies (IDT). An EnGen Cas9 sgRNA kit (NEB) was then used with each oligo to generate sgRNA by *in vitro* transcription, following the kit instructions. sgRNA were purified by phenol/chloroform extraction and ammonium acetate and ethanol precipitation, resuspended in 30 μL pyrogen free water and stored at -70 °C in aliquots of 3 μL. Fertilised embryos, identified by the presence of sperm entry points, were injected with Cas9-NLS (NEB) preincubated with sgRNAs as previously described (Chapman et al., 2022). Genotyping was done to confirm editing by extracting DNA from three randomly chosen embryos from each batch and for each guide at stage 18 as in Genomic DNA (1 μL) was amplified with PCR primers (Table 4) and Sanger sequenced. Editing was determined by comparing Sanger traces to sibling controls using TIDE (Brinkman et al., 2014).

**Table 3:**
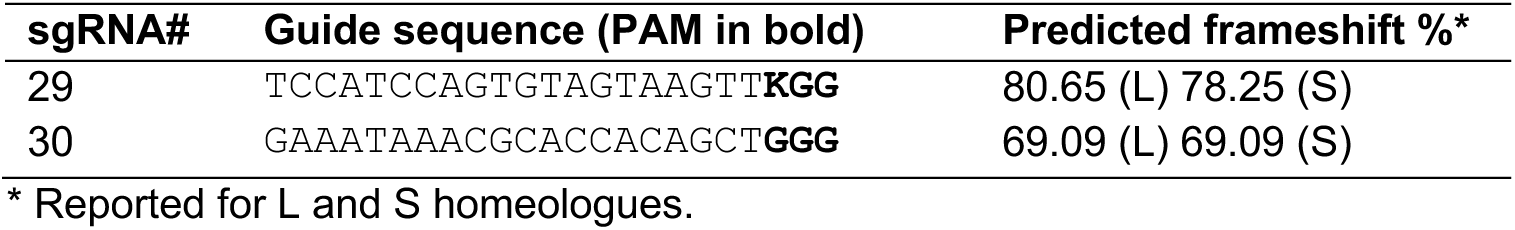
sgRNA targeting *X. laevis Tlr2*.

**Table 4:**
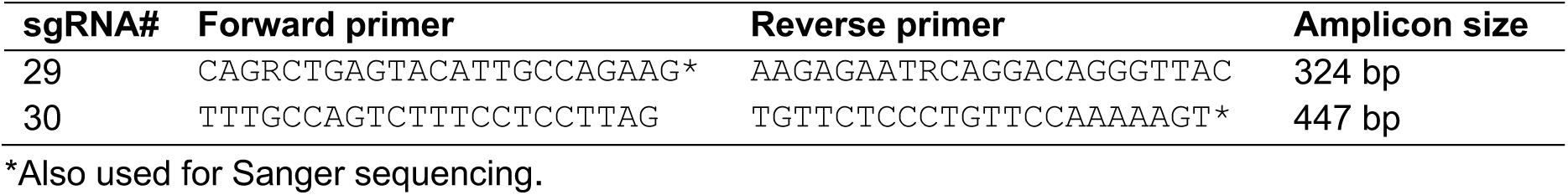
PCR amplicons for *Tlr2* CRISPR/Cas9 sequencing edits.

### LPS detection assays

Measurement of lipopolysaccharides (known as endotoxin) from samples was done using a Pierce Chromogenic Endotoxin Quant kit (Thermo Scientific), following the manufacturer’s instructions. The kit uses *Limulus* (horseshoe crab) amoebocyte lysate (LAL) assays which produce a yellow colour in proportion to the amount of LPS present. Tadpole tail tips, excised using a sterile scalpel blade, were rinsed twice in sterile 0.1 x MMR after amputation and placed in a PCR tube with 100 μL filter-sterilized NaCl/Tween solution (0.15 M NaCl, 0.1% Tween 20). Tubes were vortexed for 30 seconds and stored at -20°C until testing. Tadpole medium samples were taken directly from the 0.1 x MMR that tadpoles were swimming in. Samples were analysed in triplicate in 96 well plates with NaCl/Tween20 as the background, using a BMG Labtech FLUOstar Omega plate reader. Standard curves (high range, 0.1 to 1 endotoxin unit/mL) were prepared with each plate using a known concentration of LPS were used to convert spectrophotometric readings at 405 nm to endotoxin units (EU).

### DNA extraction and microbial sequencing

#### 16S rRNA sequencing

DNA extraction and 16S rRNA V4 hypervariable region sequencing protocols were as described by Chapman et al (2024). In short, DNA was extracted from tadpole tail clips using the DNeasy PowerLyser PowerSoil DNA extraction kit (Qiagen – Hilden, GER), and amplified and sequenced at the Argonne National Laboratory (Illinois, USA) on the Illumina MiSeq platform using primers 515F/806R (Caporaso et al., 2011) and peptide nucleic acid clamps to minimize amplification of host DNA (Lundberg et al., 2013). Reads were trimmed, quality filtered and merged using the DADA2 (v1.16) pipeline (Callahan et al., 2016). Taxonomy was assigned using the naïve Bayesian classifier method with reference to the SILVA database (v138.1) before passing to the phyloseq package (v1.42.0) (McMurdie and Holmes, 2013) in R v4.2.2 (R Development Core Team, 2023) for analysis. Sequences were aligned using DECIPHER v2.26.0 (Wright, 2015) and a maximum likelihood phylogenetic tree estimated using IQTree v2.2.2.3 (Minh et al., 2020). In total, 503 of 504 tadpole tail clip samples from 12 sibships were successfully sequenced and processed for analysis, with one sample removed from further analysis as the corresponding tadpole died and could not be scored for regeneration.

#### Whole genome (shotgun) sequencing

A total of 24 tail clippings were collected for shotgun sequencing. These tail clippings were collected from an additional spawning of a subset of three female *X. laevis* from the group that produced tadpoles for 16S rRNA sequencing (Table 5). DNA was extracted as for 16S rRNA sequencing samples. Sequencing libraries were prepared using the NEBNext Ultra II FS DNA Library Prep Kit for Illumina, using NEBNext Multiplex Oligos for Illumina (Unique Dual Index UMI Adapters DNA Set 1). Samples were sequenced at AgResearch Invermay, (Dunedin, New Zealand) on the Illumina NovaSeq platform (paired end reads, B5 and D1 – 100 bp, C1 – 150bp).

**Table 5.** Tadpole tail samples for shotgun sequencing*.

| Mother ID | Untreated | Gentamicin raised | Total # Samples |
| --- | --- | --- | --- |
| B5 | 4 (2 FR, 2 NR) | 0 | 4 |
| C1 | 6 (3 FR, 3 NR) | 6 (3 FR, 3 NR) | 12 |
| D1 | 4 (2 FR, 2 NR) | 4 (2 FR, 2 NR) | 8 |

\* Numbers in parentheses indicate the results of tail regeneration assays for tadpoles in each category. FR = Full Regeneration, NR = No Regeneration.

Reads were quality filtered using Fastp v0.23.2 (Chen, 2023) and deduplicated using a UMI deduplication script (Bossers, 2023). Host reads were identified and removed using Bowtie2 v2.4.5 (Langmead and Salzberg, 2012) with the *X. laevis* reference genome v10.1 (RefSeq GCF_017654675.1) aligned and classified using the Diamond v2.0.15 (Buchfink et al., 2021) and MEGAN v6.25.5 (Huson et al., 2016) pipeline. Read counts for each taxon were extracted from MEGAN for analysis in R.

### Statistical analyses

Unless otherwise indicated, all analyses were completed using R v4.2.2 and the Rstats package or with Graphpad PRISM v10. Categorical regeneration outcome data (FR, PG, PB and NR, as previously described) were compared using Extended Cochran– Armitage tests (ordinal chi-square: rstatix v0.7.2; (Kassambara and Mundt, 2020) with post hoc pairwise ordinal independence test (Benjamini–Hochberg correction: rcompanion v2.4.34 (Mangiafico, 2023).

### Sequencing analysis

#### Microbial 16S rRNA sequence data analysis

Reads were agglomerated at genus level for all analyses. For alpha diversity, samples were rarefied to 1750 reads after preparation of rarefaction curves (Supplementary Figure S2). The phyloseq package was used to calculate Observed and Shannon alpha diversity metrics for regeneration categories (grouped as either No Regeneration or Regeneration, the latter comprised of categories PB, PG and FR (Fig. 1b) while Faith’s phylogenetic diversity was calculated with picante v1.8.2 (Kembel et al., 2010). Significance was tested with Kruskal-Wallis and post-hoc Dunn’s tests with Bonferroni correction (FSA v0.9.4 - (Ogle et al., 2023) following Shapiro-Wilk test of normality. Boxplots were created to visualise results using ggplot2 v3.4.0 (Wickham, 2016). The relative abundance of microbial bacterial taxa in rarefied samples from the regeneration categories was plotted and compared using microshades v1.0 (Dahl et al., 2021).

For beta diversity, unrarefied samples with less than 500 reads were removed and weighted UniFrac distances calculated using the phyloseq::distance function. Hopkins Test (clustertend v1.6) (Wright et al., 2023) and a silhouette analysis (factoextra v1.0.7) (Kassambara and Mundt, 2020) were undertaken to assess clustering, and data ordinated with non-metric multidimensional scaling (NMDS) using phyloseq and ggplot2. Permutational multivariate analysis of variance (PERMANOVA – vegan v2.6-4 (adonis2 function) (Oksanen et al., 2019) was used to compare weighted UniFrac distances between regeneration categories, with Female ID and tadpole plate included as fixed effects with interactions.

Differential abundance between the two broad regeneration categories was explored using the ANCOMBC2 (v2.0.1) (Lin and Peddada, 2020) and MaAsLin2 (v1.12.0) (Mallick et al., 2021) packages. These analyses used unrarefied data and an alpha cut-off of q<0.25; taxa not seen at least 3 times in 5% of the samples were also removed. For MaAsLin2, default TSS normalisation was used. To test whether microbial taxa were consistent predictors of regeneration outcome, a random forest classification model was applied to the data using the RandomForestUtils v0.1.02 (Mallick et al., 2021) pipeline. The analysis was run with 1001 random trees, 25 repeats, and 10 cross repeats, with 20% of data held back during training for subsequent model testing.

The AMR package v2.1.1 (Berends et al., 2022) was used to assign Gram status (negative, positive or unknown) to microbial genera. Differences in the proportion of gram positive and negative reads were explored using ggplot2 and Wilcoxon’s rank sum test.

#### Microbial whole genome data

Bacterial read counts, both absolute and normalised (counts per million) were compared between regenerating and non-regenerating tadpoles, as well as between those with and without gentamicin exposure, using Wilcoxon’s rank sum test. The number of Gram-positive bacteria was also compared following assignment of Gram category with the AMR package. Before extraction of C1 sample DNA, we added 20 μl of spike in control II, low microbial yield (ZymoBIOMICS). This contains three microbes alien to the human and, to our knowledge, the *Xenopus* microbiome, in log distribution from 10^3^ to 10^5^ cells. The abundance of the three spike-in microbes was used to draw a standard curve, enabling calculation of the abundance of sample taxa as per manufacturer’s guidelines.

### Mitochondrial sequencing and analysis

Unfertilised eggs were collected from all 12 female frogs prior to embryo production for regeneration assays. Ten eggs from each female were placed into 1.5 ml microcentrifuge tubes with 100 μl of TE buffer and mechanically homogenised by pipetting. DNA was extracted from each sample by adding 10 μl of egg mixture to 150 μl Chelex-100 beads (5% suspended in TE buffer) and 1.5 μl Proteinase K and incubating at 65 °C for 4 hours, followed by 95 °C for 15 minutes. PCR to amplify the mitochondrial genome utilised universal primers from (Emser et al., 2021), with a single base substitution in the reverse primer to be specific to *X. laevis* (Fwd 5’ – TACGTGATCTGAGTTCAGACCG; Rev 5’ - GTAGGGCTTTAATCGTTGAACAAAC). Reactions consisted of 10 μl repliQa HiFi ToughMix (Quantabio - Beverly, Massachusetts, USA), 0.6 μl each of forward and reverse primer at 10 mM, 1 μl DNA template and nuclease free water added to make a total reaction volume of 20 μl. PCR comprised an initial denaturation step of 98 °C for 30 seconds, following by 35 cycles of 2 steps: 98 °C for 10 seconds / 68 °C for 2 mins 50 seconds. Final amplicons as visualised by electrophoresis on a 0.6% gel were approximately 17kb. Amplicons were purified using the NucleoSpin Gel and PCR Clean-up Kit (Macherey-Nagel, Düren, GER) as per manufacturer’s instructions, with the wash step repeated, and eluted to a final volume of 20 μl. Libraries were prepared using the NEBNext Ultra II FS DNA Library Prep Kit for Illumina, using NEBNext Multiplex Oligos for Illumina (Unique Dual Index UMI Adapters DNA Set 1) before sequencing on the Illumina MiSeq platform (150bp paired end reads) at the Otago Genomics Facility (University of Otago, New Zealand).

Sequences were assembled against the *X. laevis* mitochondrial reference genome (GenBank accession NC_001573) using Bowtie2. Areas with low sequencing coverage areas were trimmed, leaving a total of 17474 positions for analysis. Sequences were aligned using MUSCLE v5.1 (Edgar, 2004) and a maximum likelihood phylogeny constructed using IQTree with automatic model selection (GTR+I+G) by ModelFinder (Kalyaanamoorthy et al., 2017) and 1000 ultrafast bootstrap replicates (Hoang et al., 2018). The tree was visualised in R alongside offspring regeneration data using the ggtree package v3.6.2 (Hoang et al., 2018). Sequences are available at GenBank as accessions PP467366-PP467377.

## Acknowledgements

We thank Nikita Woodhead for excellent care of the *Xenopus* colony, the GAS sequencing facility at the University of Otago for Sanger sequencing, Argonne National Laboratories USA for 16S amplicon sequencing and AgResearch Invermay New Zealand for whole genome sequencing.

## Author contributions

Phoebe A. Chapman (Data curation, Formal analysis, Methodology, Visualization, Writing – original draft, review & editing)

Robert C Day (Methodology, supervision)

Daniel Hudson (Investigation, Data curation)

Joanna Ward (Methodology, Investigation, Formal analysis),

Xochitl C. Morgan (Conceptualisation, Funding acquisition, Methodology, Project administration, Resources, Supervision, Writing – review & editing)

Caroline W. Beck (Conceptualisation, Data curation, Funding acquisition, Investigation, Methodology, Formal analysis, Project administration, Resources, Supervision, Visualization, Writing – original draft, review & editing)

## Data and resource availability

16S amplicon and whole genome sequence data can be found in the SRA database (NCBI) Bioproject PRJNA823390 (https://www.ncbi.nlm.nih.gov/bioproject/?term=PRJNA823390). As the samples analysed in this manuscript are a subset of a larger BioProject, please filter by release date 2024-12-31. *Xenopus laevis* female mitogenomes can be found in GenBank as PP467366-PP467377. Code used in analyses can be found at https://gitlab.com/pachapman/xenopus_regeneration

## Supplementary data

Supporting information including five supplemental figures, 32 supplemental tables (in a single PDF) and one excel file are hosted on Figshare https://doi.org/10.6084/m9.figshare.28982285.

