## Supplemental figures and tables for "Commensal skin bacteria interact with the innate immune system to promote tail regeneration in *Xenopus laevis* tadpoles"

### Supplementary Figures and Tables

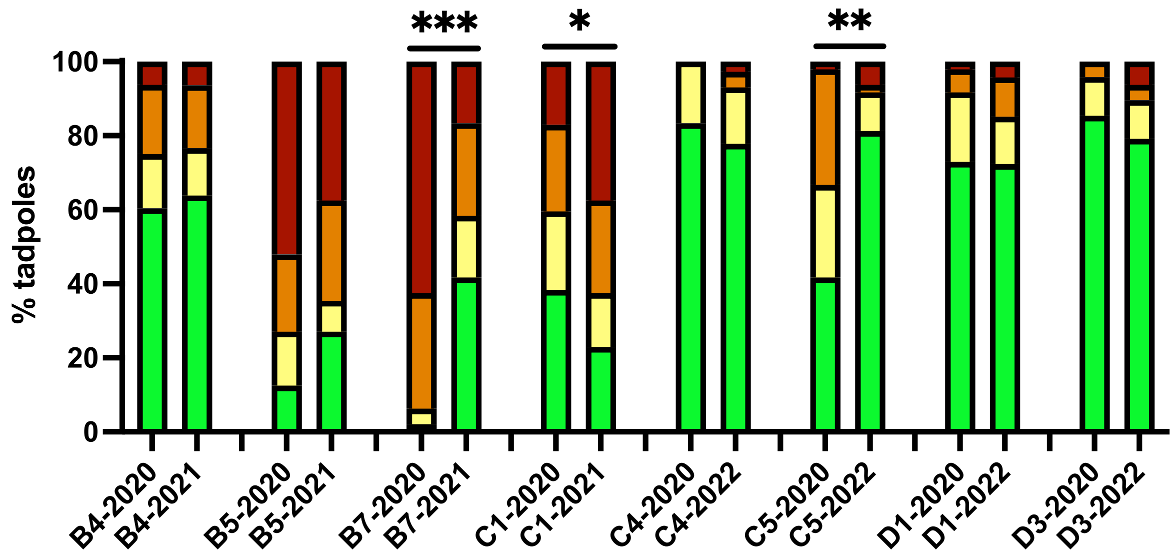

**Figure S1**. **Reproducibility of regeneration in sibships from the same female frog.**
A second spawning was obtained from 8 of the 12 females in the study. Regeneration assays were used to test for reproducibility of the regenerative outcomes from subsequent spawnings (top bar graph). For five of the eight frogs, regenerative outcomes were not significantly different between spawnings. B7, C1 and C5 females however yielded significantly different results, suggesting some impact from the environment had altered the two outcomes. N=48 for all cohorts except N=47 for B4 2021, C1 2020 and D1 2022, N=44 for C4 2020 and N=72 for C4 2022). Four frogs did not produce enough eggs for analysis. Ordinal $\chi^2$ was used to indicate significant differences in regeneration within each sibship, adjusted *p* values * *p* <0.05, ** *p*<0.01, *** *p.*<0.001, non-significant comparisons are not shown. Raw counts and analysis can be found in Table S29.

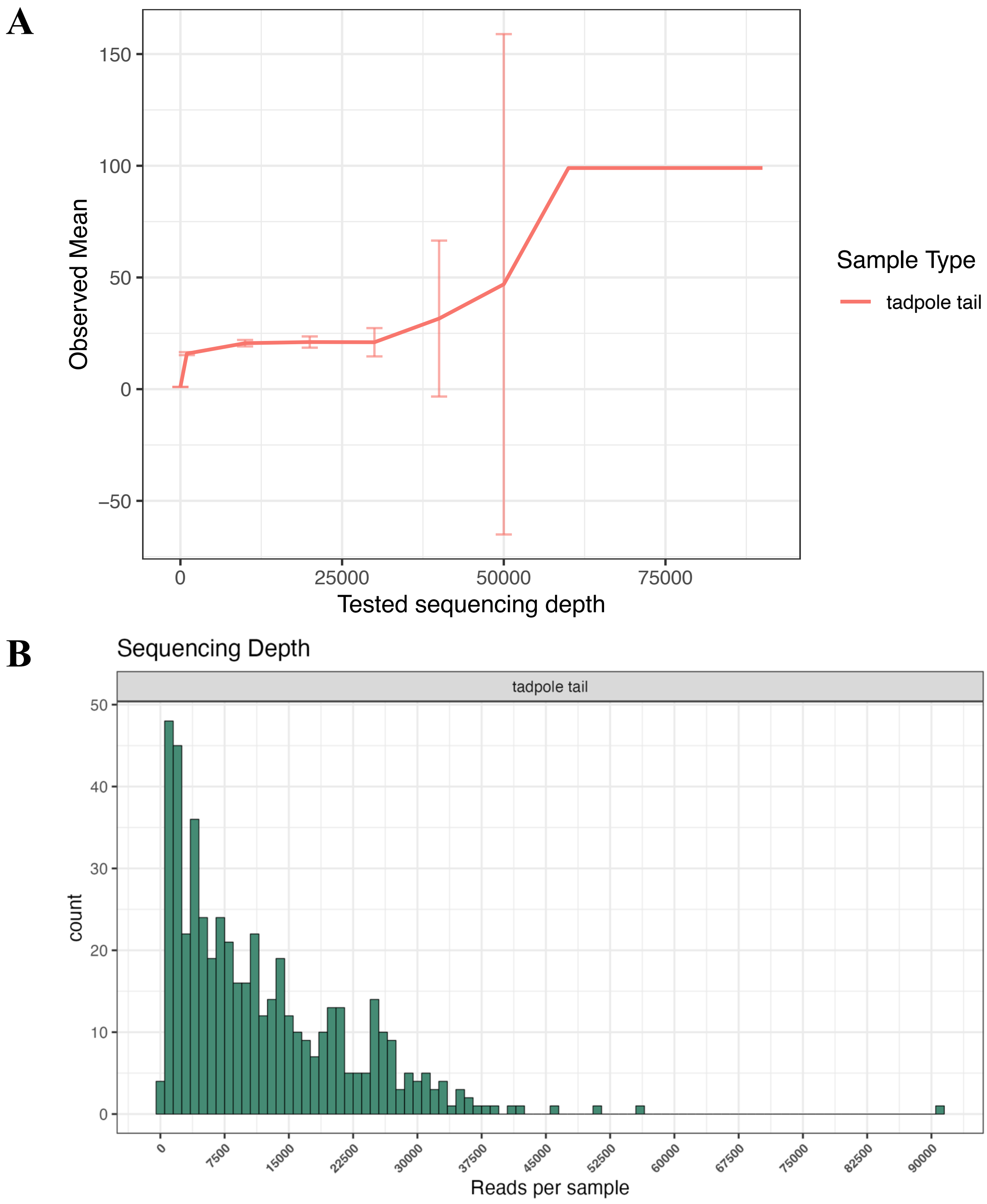

**Figure S2**. A. Rarefaction curve for 16S rRNA sequenced tadpole tail samples. B. Read count histogram.

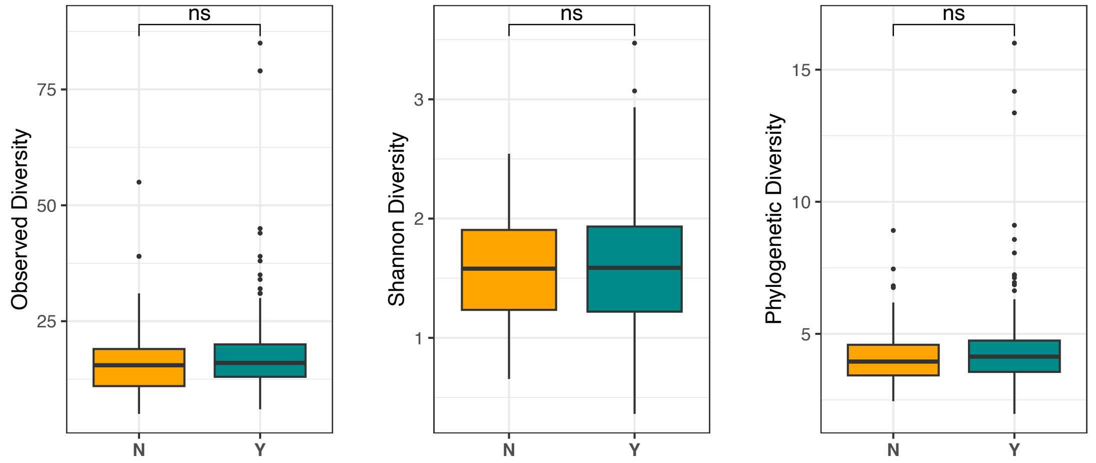

**Figure S3.** Alpha diversity of the microbial communities found on tadpole tail skin is not distinct between regenerating (Y=PB, PG, FR) and non-regenerating (N=NR) groups. Statistical analyses can be found in Table S9.

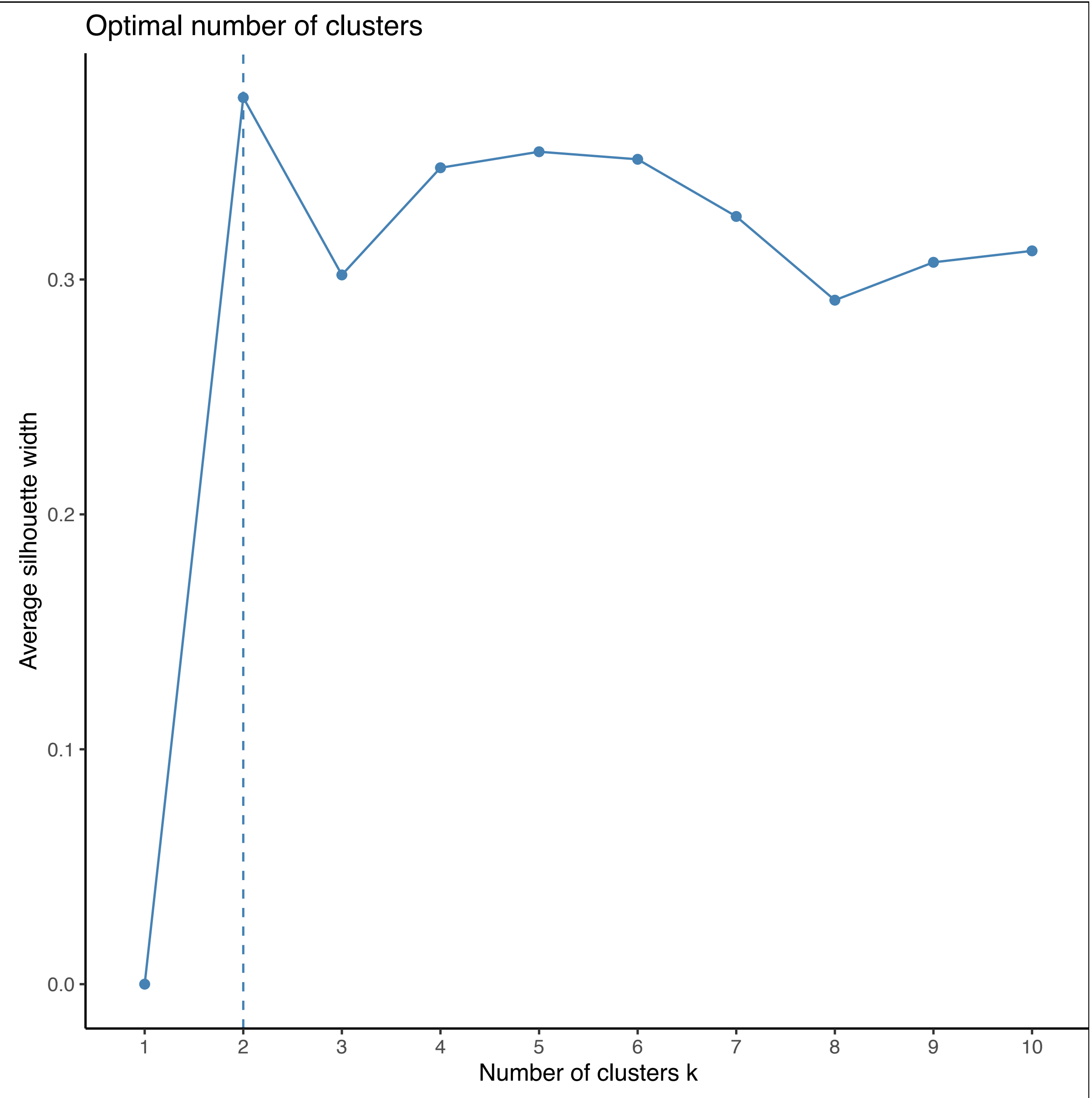

**Figure S4**. Silhouette analysis of 16S rRNA microbiome community data from tadpole tails. Hopkins statistic 0.91.

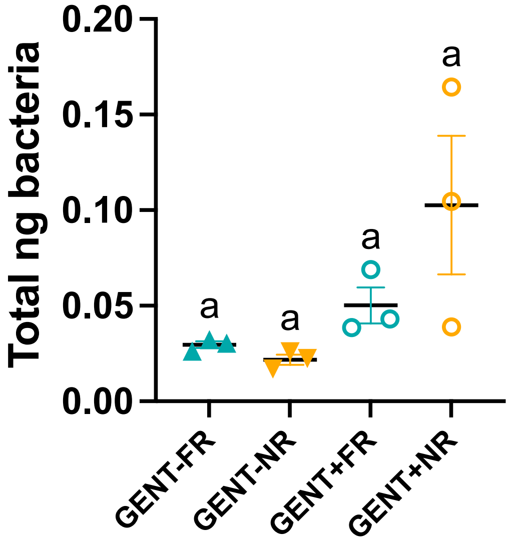

**Figure S5.** Spike-in of three commercial non-commensal standards to C1 shotgun tail genome samples before DNA isolation was used to quantify the amount of bacterial DNA in each sample. Scatterplot, 1-way ANOVA with Tukey's post hoc tests, groups that share a lower case letter are not significantly different from each other. Raw data and analysis can be found in Tables S30-S31

**Table S1: Tlr2 CRISPR/Cas9 knockdown regeneration category counts**

| | | Regeneration counts | | | | | |
|---|---|---|---|---|---|---|---|
| Treatment | | **NR** | **PB** | **PG** | **FR** | **Sample size (N)** | **% any regeneration** |
| Left sibship | Controls Cas9 | 3 | 2 | 1 | 25 | 31 | 90.3 |
| | *tlr2* sgRNA rnk29 | 17 | 3 | 1 | 1 | 22 | 22.7 |
| | *tlr2* sgRNA rnk30 | 13 | 5 | 4 | 4 | 26 | 50.0 |
| Right sibship | Controls Cas9 | 12 | 0 | 0 | 24 | 36 | 66.7 |
| | *tlr2* sgRNA rnk29 | 33 | 5 | 1 | 1 | 40 | 17.5 |
| | *tlr2* sgRNA rnk30 | 16 | 4 | 1 | 9 | 30 | 46.7 |

**Table S2: Tlr2 CRISPR/Cas9 knockdown regeneration asymptotic generalised Pearson Chi-squared test, data ordered logistically.**

| **Group** | **Chi-squared** | **Comparison** | **p.value** | **p.adjust** | **significance** |
|---|---|---|---|---|---|
| Left sibship | chi-squared = 31.033, df = 2, p-value = 0.0000001825 | control : sgRNArnk29 | 2.28E-08 | 6.84E-08 | **** |
| | | control: sgRNArnk30 | 0.00000549 | 0.00000824 | **** |
| | | sgRNArnk29: sgRNArnk30 | 0.0438 | 0.0438 | * |
| Right sibship | chi-squared = 67.464, df = 2, p-value = 0.0000000000000222 | control : sgRNArnk29 | 3.69E-08 | 0.000000111 | **** |
| | | control: sgRNArnk30 | 0.0118 | 0.0118 | * |
| | | sgRNArnk29: sgRNArnk30 | 0.00104 | 0.00156 | ** |

**Table S3: CRISPR/Cas9 editing of Tlr2 from randomly chosen embryos**

| Tadpoles | Guide RNA_sample | Editing % | | |
|---|---|---|---|---|
| | | Total | mean | SEM |
| **Left sibship** | *tlr2* sgRNA rnk29_1 | 28.6 | | |
| | *tlr2* sgRNA rnk29_2 | 53.6 | | |
| | *tlr2* sgRNA rnk29_3 | 49.1 | 43.8 | 7.69 |
| **Right sibship** | *tlr2* sgRNA rnk29_1 | 47.3 | | |
| | *tlr2* sgRNA rnk29_2 | 50.1 | | |
| | *tlr2* sgRNA rnk29_3 | 52.5 | 50.0 | 1.50 |
| **Left sibship** | *tlr2* sgRNA rnk30_1 | 6.3 | | |
| | *tlr2* sgRNA rnk30_2 | 12 | | |
| | *tlr2* sgRNA rnk30_3 | 11.5 | 9.9 | 1.82 |
| **Right sibship** | *tlr2* sgRNA rnk29_1 | 12.5 | | |
| | *tlr2* sgRNA rnk29_2 | 20.9 | | |
| | *tlr2* sgRNA rnk29_3 | 24.1 | 19.2 | 2.01 |

**Table S4: Peptidoglycan and Lipopolysaccharide treatments, category counts**

| Treatment | | Regeneration counts | | | | Sample size (N) | % any regeneration |
|---|---|---|---|---|---|---|---|
| | | NR | PB | PG | FR | | |
| **Native MMR** | none | 7 | 5 | 5 | 6 | 23 | 69.6 |
| | PG | 7 | 8 | 6 | 2 | 23 | 69.6 |
| | LPS | 8 | 9 | 1 | 4 | 22 | 63.6 |
| | PG and LPS | 3 | 5 | 4 | 11 | 23 | 87.0 |
| **Gentamicin** | none | 14 | 6 | 3 | 2 | 25 | 44.0 |
| **50 ug/ml** | PG | 7 | 9 | 3 | 5 | 24 | 70.8 |
| **MMR** | LPS | 8 | 7 | 3 | 8 | 26 | 69.2 |
| | PG and LPS | 4 | 5 | 1 | 10 | 20 | 80.0 |

**Table S5: Peptidoglycan and Lipopolysaccharide treatments regeneration asymptotic generalised Pearson Chi-squared test, data ordered logistically.**

| Group | Chi-squared | Comparison | p.value | p.adjust | significance |
|---|---|---|---|---|---|
| Control MMR | chi-squared = 9.8291, df = 3, p-value = 0.02008 | none : PG | 0.338 | 0.406 | ns |
| | | none : LPS | 0.251 | 0.376 | ns |
| | | none : PG and LPS | 0.102 | 0.204 | ns |
| | | PG : LPS | 0.777 | 0.777 | ns |
| | | PG : PG and LPS | 0.00824 | 0.0247 | * |
| | | LPS : PG and LPS | 0.00691 | 0.0247 | * |
| Gentamicin 50 ug/ml In MMR | chi-squared = 10.453, df = 3, p-value = 0.01508 | none : PG | 0.08 | 0.146 | ns |
| | | none : LPS | 0.0293 | 0.0879 | ns |
| | | none : PG and LPS | 0.00215 | 0.0129 | * |
| | | PG : LPS | 0.598 | 0.598 | ns |
| | | PG : PG and LPS | 0.0976 | 0.146 | ns |
| | | LPS : PG and LPS | 0.248 | 0.298 | ns |

**Table S6 Regeneration category counts for samples used in 16S rRNA amplicon sequencing, by mother and culture plate and pairwise comparisons of plate replicates.**

| Female-plate replicate | Regeneration counts | | | | Sample size (N) | % any regeneration | Asymptotic Linear-by-Linear Association Test | | |
|---|---|---|---|---|---|---|---|---|---|
| | FR | PG | PB | NR | | | Z | p-value | significance |
| B4-1 | 10 | 2 | 7 | 2 | 21 | 90.5 | 2.1056 | 0.0352 | * |
| B4-2 | 14 | 5 | 2 | 0 | 21 | 100.0 | | | |
| B5-1 | 2 | 2 | 8 | 9 | 21 | 57.1 | -0.4799 | 0.6313 | ns |
| B5-2 | 1 | 5 | 2 | 13 | 21 | 38.1 | | | |
| B7-1 | 1 | 1 | 8 | 11 | 21 | 47.6 | -1.1043 | 0.2695 | ns |
| B7-2 | 0 | 1 | 6 | 14 | 21 | 33.3 | | | |
| B9-1 | 20 | 1 | 0 | 0 | 21 | 100.0 | -1.1854 | 0.2359 | ns |
| B9-2 | 18 | 2 | 1 | 0 | 21 | 100.0 | | | |
| C1-1 | 10 | 4 | 2 | 4 | 20 | 69.6 | -1.3862 | 0.1657 | ns |
| C1-2 | 4 | 6 | 8 | 3 | 21 | 75.0 | | | |
| C3-1 | 18 | 1 | 1 | 1 | 21 | 83.3 | -1.5344 | 0.1249 | ns |
| C3-2 | 14 | 1 | 3 | 3 | 21 | 75.0 | | | |
| C4-1 | 16 | 5 | 0 | 0 | 21 | 87.5 | 0.7859 | 0.4319 | ns |
| C4-2 | 18 | 3 | 0 | 0 | 21 | 87.5 | | | |
| C5-1 | 11 | 5 | 5 | 0 | 21 | 87.5 | -1.3404 | 0.1801 | ns |
| C5-2 | 8 | 4 | 8 | 1 | 21 | 83.3 | | | |
| D1-1 | 18 | 2 | 0 | 1 | 21 | 83.3 | -1.2820 | 0.1998 | ns |
| D1-2 | 13 | 5 | 3 | 0 | 21 | 87.5 | | | |
| D2-1 | 18 | 2 | 1 | 0 | 21 | 87.5 | -0.5420 | 0.5879 | ns |
| D2-2 | 17 | 2 | 2 | 0 | 21 | 87.5 | | | |
| D3-1 | 18 | 2 | 1 | 0 | 21 | 87.5 | 0.3573 | 0.7209 | ns |
| D3-2 | 18 | 3 | 0 | 0 | 21 | 87.5 | | | |
| D4-1 | 13 | 3 | 4 | 1 | 21 | 83.3 | 0.5136 | 0.6075 | ns |
| D4-2 | 15 | 1 | 5 | 0 | 21 | 87.5 | | | |

**Table S7: Top five Genera for each female tank**

| Tank housing (mother) | Genus | Phylum | Mean_abundance |
|---|---|---|---|
| **Tank B B4, B5, B7, B9** | Rhizobium Genera | Proteobacteria | 749.23 |
| | Chryseobacterium | Bacteroidota | 385.07 |
| | Delftia | Proteobacteria | 67.14 |
| | Lachnospir. NK4A136 | Firmicutes | 66.60 |
| | Fluviicola | Bacteroidota | 52.86 |
| **Tank C C1, C3, C4, C5** | Aeromonas | Proteobacteria | 575.04 |
| | Rhizobium Genera | Proteobacteria | 258.20 |
| | Fluviicola | Bacteroidota | 221.07 |
| | Chryseobacterium | Bacteroidota | 161.75 |
| | Acinetobacter | Proteobacteria | 129.04 |
| **Tank D D1, D2, D3, D4** | Rhizobium Genera | Proteobacteria | 759.12 |
| | Lactiplantibacillus | Firmicutes | 254.74 |
| | Chryseobacterium | Bacteroidota | 235.67 |
| | Lachnospir. NK4A136 | Firmicutes | 67.16 |
| | Pseudomonas | Proteobacteria | 30.94 |

**Table S8 Top five genera found in each regeneration category**

| Regeneration category | Genus | Phylum | Mean_abundance |
|---|---|---|---|
| FR | Rhizobium Genera | Proteobacteria | 582.98 |
| | Chryseobacterium | Bacteroidota | 254.53 |
| | Aeromonas | Proteobacteria | 218.99 |
| | Lactiplantibacillus | Firmicutes | 111.63 |
| | Fluviicola | Bacteroidota | 83.24 |
| PG | Rhizobium Genera | Proteobacteria | 573.45 |
| | Aeromonas | Proteobacteria | 253.23 |
| | Chryseobacterium | Bacteroidota | 203.58 |
| | Fluviicola | Bacteroidota | 122.87 |
| | Lactiplantibacillus | Firmicutes | 105.42 |
| PB | Rhizobium Genera | Proteobacteria | 518.03 |
| | Aeromonas | Proteobacteria | 297.03 |
| | Chryseobacterium | Bacteroidota | 229.92 |
| | Fluviicola | Bacteroidota | 157.49 |
| | Lachnospir. NK4A136 | Firmicutes | 75.54 |
| NR | Rhizobium Genera | Proteobacteria | 579.75 |
| | Chryseobacterium | Bacteroidota | 339.59 |
| | Aeromonas | Proteobacteria | 167.29 |
| | Fluviicola | Bacteroidota | 126.13 |
| | Delftia | Proteobacteria | 121.95 |

**Table S9 Alpha diversity measures: Regenerators (PB, PG, FR categories) vs non-regenerators (NR) (Figures 3C and S4)**

| | Observed | Shannon | Phylogenetic Diversity |
|---|---|---|---|
| Shapiro-Wilk normality test W | 0.727 | 0.992 | 0.75 |
| Shapiro-Wilk normality test p-val | p<0.00001 | p=0.013 | p<0.00001 |
| Normal? | No | No | No |
| Wilcoxon rank sum test with continuity correction W | 9284 | 10470 | 9475 |
| Wilcoxon rank sum test with continuity correction p-val | 0.110 | 0.799 | 0.168 |
| Significant difference? | No | No | No |

**Table S10 NMDA PERMANOVA-Weighted UniFrac any regeneration vs. NR**

| | Df | SumOfSqs | R2 | F | Pr(>F) | % effect |
|---|---|---|---|---|---|---|
| RegenYN | 1 | 0.366 | 0.006 | 5.617 | 0.001 | 0.6 |
| FemCode | 11 | 22.179 | 0.352 | 30.973 | 0.001 | 35.2 |
| FemCode:TadpolePlate | 12 | 9.576 | 0.152 | 12.259 | 0.001 | 15.2 |
| Residual | 474 | 30.856 | 0.490 | NA | NA | 49.0 |
| Total | 498 | 62.977 | 1.000 | NA | NA | 100.0 |

**Table S11 NMDA PERMANOVA-Weighted UniFrac by mother tank (B, C or D)**

| | Df | SumOfSqs | R2 | F | Pr(>F) | % effect |
|---|---|---|---|---|---|---|
| RegenYN | 1 | 0.366 | 0.006 | 4.395 | 0.001 | 0.6 |
| Tank | 2 | 11.551 | 0.183 | 69.424 | 0.001 | 18.3 |
| Tank:FemCode | 9 | 10.628 | 0.169 | 14.194 | 0.001 | 16.9 |
| Residual | 486 | 40.432 | 0.642 | NA | NA | 64.2 |
| Total | 498 | 62.977 | 1 | NA | NA | 100.0 |

**Table S12 Random Forest analysis. Top 20 most important taxa for classifying tadpoles into regenerator or non-regenerator categories.**

| Genus | p_value | statistic | significance | Mean | SD | Min | Max | RegenYN |
|---|---|---|---|---|---|---|---|---|
| Delftia | 0.0001 | 16132 | *** | 0.0072 | 0.0030 | 0.0032 | 0.0168 | N |
| Aeromonas | 0.5524 | 11887 | ns | 0.0030 | 0.0008 | 0.0016 | 0.0048 | Y |
| Rhizobium Genera | 0.9394 | 12374 | ns | 0.0019 | 0.0005 | 0.0009 | 0.0029 | Y |
| Fluviicola | 0.3922 | 13232 | ns | 0.0019 | 0.0007 | 0.0006 | 0.0034 | N |
| Chryseobacterium | 0.0189 | 14782 | * | 0.0016 | 0.0007 | 0.0004 | 0.0029 | N |
| Bosea | 0.0060 | 9742 | ** | 0.0014 | 0.0006 | 0.0003 | 0.0027 | Y |
| Staphylococcus | 0.1956 | 13504 | ns | 0.0008 | 0.0005 | 0.0002 | 0.0019 | Y |
| Lactiplantibacillus | 0.0000 | 8944 | *** | 0.0008 | 0.0005 | 0.0001 | 0.0018 | Y |
| Undibacterium | 0.4725 | 12943 | ns | 0.0008 | 0.0004 | 0.0002 | 0.0016 | Y |
| Klebsiella | 0.0011 | 14763 | ** | 0.0008 | 0.0005 | 0.0000 | 0.0021 | N |
| Lactobacillus | 0.2365 | 13289 | ns | 0.0008 | 0.0003 | 0.0003 | 0.0014 | N |
| Lachnospir. NK4A136 | 0.7951 | 12249 | ns | 0.0005 | 0.0002 | 0.0001 | 0.0009 | N |
| Ralstonia | 0.1097 | 10864 | ns | 0.0005 | 0.0007 | -0.0008 | 0.0019 | Y |
| Vogesella | 0.1302 | 11265 | ns | 0.0004 | 0.0003 | 0.0000 | 0.0012 | Y |
| Flavobacterium | 0.3260 | 13364 | ns | 0.0004 | 0.0004 | -0.0004 | 0.0010 | Y |
| Lachnospiraceae UCG-001 | 0.2929 | 13045 | ns | 0.0004 | 0.0003 | -0.0001 | 0.0010 | N |
| Acinetobacter | 0.0641 | 10835 | ns | 0.0004 | 0.0005 | -0.0003 | 0.0017 | Y |
| Mycobacterium | 0.0071 | 13335 | ** | 0.0004 | 0.0003 | -0.0001 | 0.0010 | N |
| Oscillibacter | 0.1613 | 13121 | ns | 0.0003 | 0.0003 | -0.0001 | 0.0011 | N |
| Pedobacter | 0.8375 | 12310 | ns | 0.0003 | 0.0003 | -0.0003 | 0.0011 | Y |

**Table S13 Differential Abundance analyses with tadpole mother identity (sibship) added as a fixed effect for two comparisons.**

| | | ANCOMBC2 | | | MaAsLin2 | | |
|---|---|---|---|---|---|---|---|
| **Comparison** | **Genus** | ***p*** **val** | **q val** | **Higher in:** | ***p*** **val** | **q val** | **Higher in:** |
| **Regenerating Vs. Non-Regenerating, N=503 samples** | *Fluviicola* | ns | ns | - | 0.066 | 0.198 | Regeneration |
| | *Variovorax* | ns | ns | - | 0.040 | 0.139 | Regeneration |
| | *Herbaspirillum* | ns | ns | - | 0.065 | 0.194 | Regeneration |
| | *Streptococcus* | ns | ns | - | 0.096 | 0.243 | Non-regeneration |
| | *Klebsiella* | 0.004 | 0.221 | Non-regeneration | 0.002 | 0.010 | Non-regeneration |
| **FR vs. NR N=353 samples** | *Shinella* | ns | ns | - | 0.040 | 0.164 | FR |

* cut-off FDR q<0.25

**Table S14 Lactiplantibacillus Differential Abundance (no regeneration vs. full regeneration).**

| **Shapiro-Wilk test** | **NR** | **FR** |
|---|---|---|
| W | 0.144 | 0.4098 |
| P value | <0.0001 | <0.0001 |
| Passed normality test (alpha=0.05)? | No | No |
| P value summary | **** | **** |
| **Descriptive stats** | **NR** | **FR** |
| Number of values | 56 | 259 |
| Minimum | 0 | 0 |
| Maximum | 0.04971 | 0.92 |
| Range | 0.04971 | 0.92 |
| Mean | 0.00104 | 0.06379 |
| Std. Deviation | 0.00665 | 0.17890 |
| Std. Error of Mean | 0.00089 | 0.01112 |
| **Mann Whitney test** | | **NR vs. FR** |
| P value | | <0.0001 |
| Exact or approximate P value? | | Exact |
| P value summary | | **** |
| Significantly different (P < 0.05)? | | Yes |
| One- or two-tailed P value? | | Two-tailed |
| Sum of ranks in column A,B | | 6426 , 43345 |
| Mann-Whitney U | | 4830 |

**Table S15 Pseudomonas Differential Abundance (no regeneration vs. full regeneration).**

| **Shapiro-Wilk test** | **NR** | **FR** |
|---|---|---|
| W | 0.4028 | 0.63 |
| P value | <0.0001 | <0.0001 |
| Passed normality test (alpha=0.05)? | No | No |
| P value summary | **** | **** |
| **Descriptive stats** | **NR** | **FR** |
| Number of values | 56 | 259 |
| Minimum | 0 | 0 |
| Maximum | 0.188 | 0.2183 |
| Range | 0.188 | 0.2183 |
| Mean | 0.01035 | 0.01611 |
| Std. Deviation | 0.02841 | 0.02714 |
| Std. Error of Mean | 0.00380 | 0.00169 |
| **Mann Whitney test** | **NR vs. FR** | |
| P value | | 0.0058 |
| Exact or approximate P value? | | Exact |
| P value summary | | ** |
| Significantly different (P < 0.05)? | | Yes |
| One- or two-tailed P value? | | Two-tailed |
| Sum of ranks in column A,B | | 7195 , 42575 |
| Mann-Whitney U | | 5599 |

**Table S16 Sphingobium Differential Abundance (no regeneration vs. full regeneration).**

| **Shapiro-Wilk test** | **NR** | **FR** |
|---|---|---|
| W | 0.2784 | 0.5511 |
| P value | <0.0001 | <0.0001 |
| Passed normality test (alpha=0.05)? | No | No |
| P value summary | **** | **** |
| **Descriptive stats** | **NR** | **FR** |
| Number of values | 56 | 259 |
| Minimum | 0 | 0 |
| Maximum | 0.048 | 0.06 |
| Range | 0.048 | 0.06 |
| Mean | 0.0021 | 0.0054 |
| Std. Deviation | 0.0085 | 0.0113 |
| Std. Error of Mean | 0.0011 | 0.0007 |
| **Mann Whitney test** | | **NR vs. FR** |
| P value | | 0.0013 |
| Exact or approximate P value? | | Exact |
| P value summary | | ** |
| Significantly different (P < 0.05)? | | Yes |
| One- or two-tailed P value? | | Two-tailed |
| Sum of ranks in column A,B | | 7258 , 42512 |
| Mann-Whitney U | | 5662 |

**Table S17 Cupriavidus Differential Abundance (no regeneration vs. full regeneration**

| **Shapiro-Wilk test** | **NR** | **FR** |
|---|---|---|
| W | 0.1439 | 0.2486 |
| P value | <0.0001 | <0.0001 |
| Passed normality test (alpha=0.05)? | No | No |
| P value summary | **** | **** |
| **Descriptive stats** | **NR** | **FR** |
| Number of values | 56 | 259 |
| Minimum | 0 | 0 |
| Maximum | 0.00685 | 0.07943 |
| Range | 0.00685 | 0.07943 |
| Mean | 0.00014 | 0.00193 |
| Std. Deviation | 0.00093 | 0.00823 |
| Std. Error of Mean | 0.00012 | 0.00051 |
| **Mann Whitney test** | | **NR vs FR** |
| P value | | 0.0129 |
| Exact or approximate P value? | | Exact |
| P value summary | | * |
| Significantly different (P < 0.05)? | | Yes |
| One- or two-tailed P value? | | Two-tailed |
| Sum of ranks in column A,B | | 7950 , 41821 |
| Mann-Whitney U | | 6354 |

**Table S18 Klebsiella Differential Abundance (no regeneration vs. full regeneration)**

| **Shapiro-Wilk test** | **NR** | **FR** |
|---|---|---|
| W | 0.4619 | 0.2982 |
| P value | <0.0001 | <0.0001 |
| Passed normality test (alpha=0.05)? | No | No |
| P value summary | **** | **** |
| **Descriptive stats** | **NR** | **FR** |
| Number of values | 56 | 259 |
| Minimum | 0 | 0 |
| Maximum | 0.07657 | 0.1006 |
| Range | 0.07657 | 0.1006 |
| Mean | 0.00545 | 0.00285 |
| Std. Deviation | 0.01338 | 0.01054 |
| Std. Error of Mean | 0.00179 | 0.00065 |
| **Mann Whitney test** | | **NR vs FR** |
| P value | | 0.0012 |
| Exact or approximate P value? | | Exact |
| P value summary | | ** |
| Significantly different (P < 0.05)? | | Yes |
| One- or two-tailed P value? | | Two-tailed |
| Sum of ranks in column A,B | | 10316, 39454 |
| Mann-Whitney U | | 5784 |

**Table S19 Delftia Differential Abundance (no regeneration vs. full regeneration)**

| **Shapiro-Wilk test** | **NR** | **FR** |
|---|---|---|
| W | 0.7795 | 0.5448 |
| P value | <0.0001 | <0.0001 |
| Passed normality test (alpha=0.05)? | No | No |
| P value summary | **** | **** |
| **Descriptive stats** | **NR** | **FR** |
| Number of values | 56 | 259 |
| Minimum | 0 | 0 |
| Maximum | 0.3017 | 0.1949 |
| Range | 0.3017 | 0.1949 |
| Mean | 0.06968 | 0.01882 |
| Std. Deviation | 0.0873 | 0.0382 |
| Std. Error of Mean | 0.01167 | 0.002373 |
| **Mann Whitney test** | | **NR vs FR** |
| P value | | <0.0001 |
| Exact or approximate P value? | | Exact |
| P value summary | | **** |
| Significantly different (P < 0.05)? | | Yes |
| One- or two-tailed P value? | | Two-tailed |
| Sum of ranks in column A,B | | 11721, 38049 |
| Mann-Whitney U | | 4379 |

**Table S20 LAL assay results for tails (Endotoxin Units/mL) (Figure 6A)**

| Reg_Y Gent_N | Reg_N Gent_N | Reg_Y Gent_Y | Reg_N Gent Y |
|---|---|---|---|
| 3.226 | 2.989 | -0.098 | 0.126 |
| 2.785 | 1.580 | -0.246 | 0.052 |
| 2.79 | 1.884 | -0.224 | -0.282 |
| 0.731 | 1.962 | 0.021 | 0.003 |
| 0.65 | 2.574 | -0.038 | -0.052 |
| 1.337 | 0.643 | -0.074 | -0.029 |
| 0.155 | 0.22 | 0.337 | 0.450 |
| 0.083 | 1.171 | 0.428 | 0.096 |
| 0.671 | 0.113 | 0.236 | 0.389 |

**Table S21 Statistical comparison of LAL assay results for tails by 1-way ANOVA (Figure 6A)**

| Tukey's multiple comparisons test | Mean Diff. | Summary | P Adj. |
|---|---|---|---|
| **Reg_Y Gent_N** vs. **Reg_N Gent_N** | -0.07867 | ns | 0.9968 |
| **Reg_Y Gent_N** vs. **Reg_Y Gent_Y** | 1.343 | ** | 0.0069 |
| **Reg_Y Gent_N** vs. **Reg_N Gent Y** | 1.297 | ** | 0.0095 |
| **Reg_N Gent_N** vs. **Reg_Y Gent_Y** | 1.422 | ** | 0.0040 |
| **Reg_N Gent_N** vs. **Reg_N Gent Y** | 1.376 | ** | 0.0055 |
| **Reg_Y Gent_Y** vs. **Reg_N Gent Y** | -0.04567 | ns | 0.9994 |

Table S22 LAL assay results for tadpole media (Endotoxin Units/mL) (Figure 6B)

| | Sibship1 | | Sibship2 | |
|---|---|---|---|---|
| Media | Gent_Y | Gent_N | Gent_Y | Gent_N |
| 0.039 | 0.698 | 8.909 | 1.125 | 8.28 |
| 0.055 | 1.019 | 8.541 | 0.657 | 8.719 |

Table S22 Statistical comparison of LAL assay results by ANOVA (Figure 6B)

| Holm-Šídák's multiple comparisons test | Mean Diff. | Summary | P Adj. |
|---|---|---|---|
| Sibship 1 Gent_Y vs. Sibship1 Gent_N | -7.867 | **** | <0.0001 |
| Sibship 1 Gent_Y vs. Sibship2 Gent_Y | -0.033 | ns | 0.9147 |
| Sibship 1 Gent_Y vs. Sibship2 Gent_N | -7.641 | **** | <0.0001 |
| Sibship1 Gent_N vs. Sibship2 Gent_Y | 7.834 | **** | <0.0001 |
| Sibship1 Gent_N vs. Sibship2 Gent_N | 0.226 | ns | 0.7225 |
| Sibship2 Gent_Y vs. Sibship2 Gent_N | -7.609 | **** | <0.0001 |

**Table S24 Shotgun whole genome normalised bacterial reads (for Figure 6C**

| Row.names | total_assigned | Bacteria_nospike | Regen? | Gent? | Proportion bacterial | % bacterial |
|---|---|---|---|---|---|---|
| trim_B5_3_1_S5 | 45931261 | 105390 | N | N | 0.00229 | 0.22945 |
| trim_B5_3_2_S4* | 37033091 | 323099 | N | N | 0.00872 | 0.87246 |
| trim_B5_3_7_S3 | 36384582 | 85867 | Y | N | 0.00236 | 0.23600 |
| trim_B5_3_8_S10 | 40934212 | 44567 | Y | N | 0.00109 | 0.10887 |
| trim_D1_4_25_S9 | 40001883 | 33947 | Y | N | 0.00085 | 0.08486 |
| trim_D1_4_26_S7 | 41273491 | 49189 | Y | N | 0.00119 | 0.11918 |
| trim_D1_4_31_S2 | 45560679 | 25742 | N | N | 0.00057 | 0.05650 |
| trim_D1_4_45_S6 | 46796129 | 31100 | N | N | 0.00066 | 0.06646 |
| trim_D1_G1_39_S8 | 52404117 | 12071 | N | Y | 0.00023 | 0.02303 |
| trim_D1_G2_25_S1 | 42693469 | 25028 | N | Y | 0.00059 | 0.05862 |
| trim_D1_G2_31_S12 | 34446662 | 34389 | Y | Y | 0.00100 | 0.09983 |
| trim_D1_G2_35_S11 | 43390731 | 18301 | Y | Y | 0.00042 | 0.04218 |
| trim_XenSpike_10C1_10_G7_S10 | 112845693 | 36932 | Y | Y | 0.00033 | 0.03273 |
| trim_XenSpike_11C1_11_H7_S11 | 189201328 | 165847 | Y | Y | 0.00088 | 0.08766 |
| trim_XenSpike_12C1_12_A6_S12 | 130195965 | 42515 | N | Y | 0.00033 | 0.03265 |
| trim_XenSpike_1C1_1_F8_S1 | 129453835 | 69219 | N | N | 0.00053 | 0.05347 |
| trim_XenSpike_2C1_2_G8_S2 | 95368123 | 76359 | Y | N | 0.00080 | 0.08007 |
| trim_XenSpike_3C1_3_H8_S3 | 123677773 | 74134 | Y | N | 0.00060 | 0.05994 |
| trim_XenSpike_4C1_4_A7_S4 | 103976851 | 42654 | N | N | 0.00041 | 0.04102 |
| trim_XenSpike_5C1_5_B7_S5 | 96076610 | 99372 | Y | N | 0.00103 | 0.10343 |
| trim_XenSpike_6C1_6_C7_S6 | 91545097 | 46608 | N | N | 0.00051 | 0.05091 |
| trim_XenSpike_7C1_7_D7_S7 | 97517274 | 63826 | N | Y | 0.00065 | 0.06545 |
| trim_XenSpike_8C1_8_E7_S8 | 49819742 | 107849 | N | Y | 0.00216 | 0.21648 |
| trim_XenSpike_9C1_9_F7_S9 | 128305599 | 131744 | Y | Y | 0.00103 | 0.10268 |

* Outlier removed from analysis.

**Table S25: Statistical analysis of normalised shotgun bacterial read counts Full Regeneration (FR) vs. No Regeneration (NR), For Figure 6C.**

| **Descriptive statistics** | **NR** | **FR** |
|---|---|---|
| Number of values | 12 | 11 |
| Minimum | 0.0230 | 0.0327 |
| Maximum | 0.236 | 0.2295 |
| Range | 0.213 | 0.1967 |
| Mean | 0.0991 | 0.0784 |
| Std. Deviation | 0.0661 | 0.0555 |
| Std. Error of Mean | 0.0191 | 0.0167 |
| **Shapiro-Wilk test** | **NR** | **FR** |
| W | 0.8596 | 0.73 |
| P value | 0.0483 | 0.0011 |
| Passed normality test (alpha=0.05)? | No | No |
| P value summary | * | ** |
| **Mann Whitney test** | | **NR vs FR** |
| P value | | 0.2316 |
| Exact or approximate P value? | | Exact |
| P value summary | | ns |
| Significantly different (P < 0.05)? | | No |
| One- or two-tailed P value? | | Two-tailed |
| Sum of ranks in column A,B | | 176, 100 |
| Mann-Whitney U | | 45 |

**Table S26: Statistical analysis of normalised shotgun bacterial read counts Gentamicin raised vs. naturally raised For Figure 6D.**

| **Descriptive statistics** | **MMR** | **GENT** |
|---|---|---|
| Number of values | 13 | 10 |
| Minimum | 0.0410 | 0.0230 |
| Maximum | 0.2360 | 0.2165 |
| Range | 0.1950 | 0.1934 |
| Mean | 0.0992 | 0.0761 |
| Std. Deviation | 0.0639 | 0.0570 |
| Std. Error of Mean | 0.0177 | 0.0180 |
| Test for normal distribution | | |
| **Shapiro-Wilk test** | **MMR** | **GENT** |
| W | 0.7713 | 0.8132 |
| P value | 0.0032 | 0.0209 |
| Passed normality test (alpha=0.05)? | No | No |
| P value summary | ** | * |
| Number of values | 13 | 10 |
| **Mann Whitney test** | | **MMR vs GENT** |
| P value | | 0.2316 |
| Exact or approximate P value? | | Exact |
| P value summary | | ns |
| Significantly different (P < 0.05)? | | No |
| One- or two-tailed P value? | | Two-tailed |
| Sum of ranks in column A,B | | 176, 100 |
| Mann-Whitney U | | 45 |

**Table S27 Tail regeneration category counts, vancomycin and gentamycin treatments (Figure 8)**

| Tadpole sibship # | Treatment-plate | NR | PB | PG | FR | Sample size N |
|---|---|---|---|---|---|---|
| 1 | MMR-1 | 4 | 3 | 1 | 22 | 30 |
| | MMR-2 | 5 | 3 | 1 | 17 | 26 |
| | MMR-3 | 2 | 3 | 4 | 23 | 32 |
| | GENT-1 | 8 | 5 | 0 | 5 | 18 |
| | GENT-2 | 7 | 6 | 5 | 0 | 18 |
| | GENT-3 | 12 | 4 | 1 | 3 | 20 |
| | VANC-1 | 18 | 3 | 3 | 4 | 28 |
| | VANC-2 | 18 | 6 | 3 | 2 | 29 |
| | VANC-3 | 20 | 3 | 2 | 4 | 29 |
| 2 | MMR-1 | 8 | 3 | 1 | 15 | 27 |
| | MMR-2 | 16 | 3 | 0 | 4 | 23 |
| | MMR-3 | 9 | 8 | 1 | 7 | 25 |
| | GENT-1 | 21 | 2 | 2 | 2 | 27 |
| | GENT-2 | 14 | 3 | 7 | 3 | 27 |
| | GENT-3 | 18 | 1 | 2 | 7 | 28 |
| | VANC-1 | 19 | 3 | 4 | 4 | 30 |
| | VANC-2 | 21 | 4 | 0 | 5 | 30 |
| | VANC-3 | 18 | 4 | 3 | 4 | 29 |
| 3 | MMR-1 | 5 | 2 | 1 | 20 | 28 |
| | MMR-2 | 2 | 0 | 2 | 24 | 28 |
| | MMR-3 | 3 | 3 | 3 | 20 | 29 |
| | MMR-4 | 1 | 0 | 4 | 20 | 25 |
| | GENT-1 | 14 | 3 | 8 | 8 | 33 |
| | GENT-2 | 12 | 8 | 3 | 2 | 25 |
| | GENT-3 | 4 | 5 | 14 | 9 | 32 |
| | GENT-4 | 6 | 3 | 6 | 13 | 28 |
| | VANCO-1 | 12 | 3 | 6 | 15 | 36 |
| | VANCO-2 | 7 | 3 | 3 | 16 | 29 |
| | VANCO-3 | 8 | 1 | 3 | 19 | 31 |
| | VANCO-4 | 2 | 6 | 2 | 20 | 30 |
| 4 | MMR-1 | 17 | 4 | 1 | 8 | 30 |
| | MMR-2 | 10 | 8 | 2 | 10 | 30 |
| | MMR-3 | 12 | 7 | 3 | 10 | 32 |
| | MMR-4 | 11 | 3 | 2 | 11 | 27 |
| | GENT-1 | 20 | 4 | 3 | 1 | 28 |
| | GENT-2 | 22 | 3 | 5 | 2 | 32 |
| | GENT-3 | 23 | 1 | 4 | 3 | 31 |
| | GENT-4 | 22 | 4 | 4 | 2 | 32 |
| | VANCO-1 | 24 | 4 | 3 | 1 | 32 |
| | VANCO-2 | 22 | 3 | 2 | 8 | 35 |
| | VANCO-3 | 10 | 6 | 5 | 8 | 29 |
| | VANCO-4 | 15 | 6 | 6 | 5 | 32 |

**Table S28 Effect of raising tadpoles in antibiotics on regeneration asymptotic generalised Pearson Chi-squared test, data ordered logistically (Figure 8)**

| Sibship # | Comparison | p.value | p.adjust | Significance |
|---|---|---|---|---|
| 1 | Gentamicin vs. vancomycin | 0.197 | 0.197 | ns |
| | MMR vs. gentamicin | 6.33E-11 | 9.50E-11 | **** |
| | MMR vs. vancomycin | 4.44E-16 | 1.33E-15 | **** |
| 2 | Gentamicin vs. vancomycin | 0.723 | 0.723 | ns |
| | MMR vs. gentamicin | 0.0134 | 0.0201 | * |
| | MMR vs. vancomycin | 0.00417 | 0.0125 | * |
| 3 | Gentamicin vs. vancomycin | 0.00206 | 0.00206 | ** |
| | MMR vs. gentamicin | 0.000521 | 0.000782 | *** |
| | MMR vs. vancomycin | 1.50E-10 | 4.50E-10 | **** |
| 4 | Gentamicin vs. vancomycin | 0.00806 | 0.0121 | * |
| | MMR vs. gentamicin | 0.00000114 | 0.00000342 | **** |
| | MMR vs. vancomycin | 0.0147 | 0.0147 | * |

**Table S29 Regeneration category counts of tadpoles derived from consecutive spawns from the same female frog. Asymptotic Linear-by-Linear Association Test (Figure S1)**

| Comparison (female_year) | p.value | significance |
|---|---|---|
| B4_2020 vs. B4_2021 | 0.808 | ns |
| B5_2020 vs. B5_2021 | 0.114 | ns |
| B7_2020 vs. B7_2021 | 7.06E-09 | **** |
| C1_2020 vs. C1_2022 | 0.0173 | * |
| C4_2020 vs. C4_2022 | 0.161 | ns |
| C5_2020 vs. C5_2022 | 0.00104 | ** |
| D1_2020 vs. D1_2022 | 0.559 | ns |
| D3_2020 vs. D3_2022 | 0.182 | ns |

**Table S30 Figure S5**

| Sample | Regeneration | Gentamicin | total_ng |
|---|---|---|---|
| trim_XenSpike_10C1_10_G7_S10 | Y | Y | 0.03859159 |
| trim_XenSpike_11C1_11_H7_S11 | Y | Y | 0.04316589 |
| trim_XenSpike_12C1_12_A6_S12 | N | Y | 0.10452818 |
| trim_XenSpike_1C1_1_F8_S1 | N | N | 0.02240141 |
| trim_XenSpike_2C1_2_G8_S2 | Y | N | 0.03036359 |
| trim_XenSpike_3C1_3_H8_S3 | Y | N | 0.03226991 |
| trim_XenSpike_4C1_4_A7_S4 | N | N | 0.0258809 |
| trim_XenSpike_5C1_5_B7_S5 | Y | N | 0.0262268 |
| trim_XenSpike_6C1_6_C7_S6 | N | N | 0.01695871 |
| trim_XenSpike_7C1_7_D7_S7 | N | Y | 0.16444212 |
| trim_XenSpike_8C1_8_E7_S8 | N | Y | 0.03898403 |
| trim_XenSpike_9C1_9_F7_S9 | Y | Y | 0.06889198 |

**Table S31 Figure S5 1 way ANOVA**

| Tukey's multiple comparisons test | Mean Diff. | 95.00% CI of diff. | Summary | Adjusted P Value |
|---|---|---|---|---|
| GENT_N REG_N vs. GENT_N REG_Y | -0.007873 | -0.09294 to 0.07720 | ns | 0.9903 |
| GENT_N REG_N vs. GENT_Y REG_N | -0.0809 | -0.1660 to 0.004166 | ns | 0.0624 |
| GENT_N REG_N vs. GENT_Y REG_Y | -0.02847 | -0.1135 to 0.05660 | ns | 0.715 |
| GENT_N REG_Y vs. GENT_Y REG_N | -0.07303 | -0.1581 to 0.01204 | ns | 0.0948 |
| GENT-REG_Y vs. GENT_Y REG_Y | -0.0206 | -0.1057 to 0.06447 | ns | 0.8635 |
| GENT+REG_N v s. GENT_Y REG_Y | 0.05243 | -0.03264 to 0.1375 | ns | 0.2731 |

**Table S32 Read count summary, 503 samples 16S rRNA from tails**

| SampleType | Samples | Min | Median | Mean | Max | SD | Total |
|---|---|---|---|---|---|---|---|
| tadpole tail | 503 | 82 | 9093 | 11868.9662 | 91228 | 10525.9963 | 5970090 |
